# Molecular architecture of 40S initiation complexes on the Hepatitis C virus IRES: from ribosomal attachment to eIF5B-mediated reorientation of initiator tRNA

**DOI:** 10.1101/2021.12.10.472117

**Authors:** Zuben P. Brown, Irina S. Abaeva, Swastik De, Christopher U.T. Hellen, Tatyana V. Pestova, Joachim Frank

## Abstract

Hepatitis C virus mRNA contains an internal ribosome entry site (IRES) that mediates end-independent translation initiation, requiring a subset of eukaryotic initiation factors (eIFs). Direct binding of the IRES to the 40S subunit places the initiation codon into the P site, where it base-pairs with eIF2-bound Met-tRNA_i_^Met^ forming a 48S initiation complex. Then, eIF5 and eIF5B mediate subunit joining. Initiation can also proceed without eIF2, in which case Met-tRNA_i_^Met^ is recruited directly by eIF5B. Here, we present cryo-EM structures of IRES initiation complexes at resolutions up to 3.5 Å that cover all major stages from initial ribosomal association, through eIF2-containing 48S initiation complexes, to eIF5B-containing complexes immediately prior to subunit joining. These structures provide insights into the dynamic network of 40S/IRES contacts, highlight the role for IRES domain II, and reveal conformational changes that occur during the transition from eIF2- to eIF5B-containing 48S complexes that prepare them for subunit joining.

## INTRODUCTION

The canonical initiation process begins with formation of the 43S preinitiation complex (PIC) comprising the 40S ribosomal subunit, the eIF2•GTP/Met-tRNA_i_^Met^ ternary complex (eIF2-TC), eIF1, eIF1A and eIF3 (Jackson et al., 2010). The 43S PIC attaches to the capped 5ʹ-terminal region of mRNA and then scans to the initiation codon in a favorable nucleotide context (containing A/G and G at the -3 and +4 positions relative to the AUG, respectively) where it stops and forms the 48S initiation complex (IC) with established codon-anticodon base-pairing. Attachment is mediated by eIFs 4A, 4B and eIF4F, which cooperatively unwind the cap-proximal region allowing attachment and also assist 43S PIC scanning. eIF1, in cooperation with eIF1A, induces an ’open’ scanning-competent conformation of the 43S PIC and monitors the fidelity of initiation codon selection (Pestova et al., 1998a; Pestova and Kolupaeva, 2002; Passmore et al., 2007; Hussain et al., 2014). Establishment of codon-anticodon base-pairing in the 48S IC leads to dissociation of eIF1 and eIF5-induced hydrolysis of eIF2-bound GTP, and thereby switches the 40S subunit to the ’closed’ conformation (Unbehaun et al., 2004; Maag et al., 2005). After that, eIF5B, in its GTP-bound form, displaces residual eIF2•GDP (Pisarev et al., 2006) and promotes joining of the 60S subunit (Pestova et al., 2000). Interaction of eIF5B with eIF1A enhances eIF5B’s subunit joining activity and the hydrolysis of eIF5B-bound GTP, leading to coupled release of eIF5B•GDP and eIF1A from the assembled 80S ribosome (Marintchev et al., 2003; Acker et al., 2006; Nag et al., 2016).

A number of viral mRNAs contain internal ribosomal entry sites (IRESs), structured RNA regions that mediate cap-independent initiation of translation using a subset of the eIFs that are required by canonical initiation. All IRES-mediated initiation mechanisms are based on non-canonical interactions of IRESs with canonical components of the translation apparatus (Jackson et al, 2010). The ∼300nt-long hepatitis C virus (HCV) IRES is located in the 5ʹ-terminal region of the viral genome and epitomizes a class of related RNA elements. HCV-like IRESs occur in the genomes of pestiviruses (e.g., classical swine fever virus (CSFV)), some pegiviruses and numerous members of *Picornaviridae* (Arhab et al., 2020). The HCV IRES comprises three domains (II– IV), with domain III divided into several subdomains (Figure 1A). Ribosomal recruitment of HCV and HCV-like IRESs occurs by direct binding of the IRES to the 40S subunit and does not involve scanning, group 4 eIFs or eIF1 (Pestova et al., 1998b). Domain III binds at the back of the 40S subunit, whereas the long, bent domain II loops out and reaches into the E site. The sites of interaction with the 40S subunit include domains IIIa and IIIc (which bind to eS1, eS26 and eS27), the apex of domain IIId (which base-pairs to expansion segment (ES) 7 of 18S rRNA), domain IIIe (which interacts with helix (h) 26 of 18S rRNA), and the apex of domain II, which interacts with uS7 and eS25 in the head and uS11 on the platform of the 40S subunit, intercalating into the mRNA binding channel and causing tilting of the head and forcing the 40S subunit to adopt the open conformation (Kolupaeva et al., 2000; Kieft et al., 2001; Malygin et al., 2013a; 2013b; Hashem et al., 2013; Angulo et al., 2016; Matsuda and Mauro, 2014; Quade et al., 2015; Yamamoto et al., 2015; Yokoyama et al., 2019).

**Figure 1.**
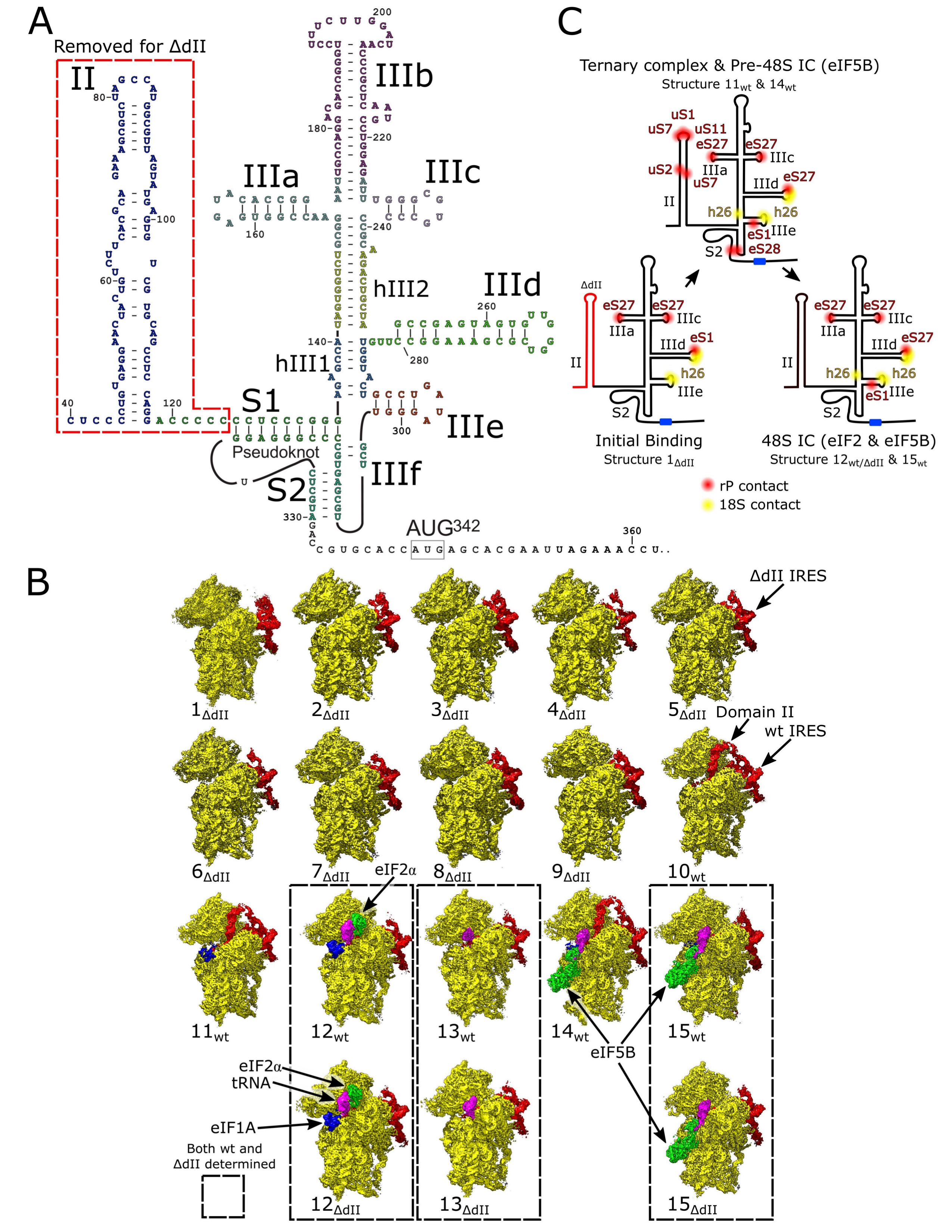
Overview of HCV IRES-mediated initiation complexes. **(A)** Secondary structure of the HCV IRES. **(B)** Segmented maps of labeled IRES ICs, showing the 40S subunit (yellow), IRES (red), eIF1A (blue), Met-tRNAi^Met^ (magenta), and initiation factors eIF2 or eIF5B (green). Complexes were assembled on the *wt* or ΔdII IRES. Complexes that share identical 40S subunit conformation and factor composition are enclosed by dashed lines. **(C)** Contacts between the IRES and 40S subunit. Ribosomal proteins (red), 18S rRNA (yellow), and the AUG codon (blue) are marked for the labeled complexes. Complete pathway in Figure S4 and Table S4. See also Figures S1-S4 and Tables S1-S4.

Although domain II does not contribute to the affinity of the HCV IRES to the 40S subunit (e.g. Kieft et al., 2001; Spahn et al., 2001), the open conformation of the 40S subunit promoted by domain II facilitates loading of the region containing the initiation codon into the mRNA binding channel, accounting for the stimulatory role of domain II during initiation on HCV-like IRESs (Honda et al., 1996; Reynolds et al., 1996; Filbin and Kieft, 2011; Odreman-Macchioli et al., 2001). Upon binding to the 40S subunit, domain IV of the HCV IRES unfolds, and the initiation codon is placed in the immediate vicinity of the P site, where it base-pairs with the anticodon of Met-tRNA_i_^Met^ as a part of the eIF2-TC, leading to formation of the 48S IC (Pestova et al., 1998b). After that, eIF5 and eIF5B mediate the subunit joining step to complete formation of the elongation-competent 80S ribosome (Locker et al., 2007; Pestova et al., 2008; Terenin et al., 2008).

Notably, in 80S complexes assembled on the HCV IRES, the P-site Met-tRNA_i_^Met^ and eIF5B•GTP, which correspond to the last stage in the initiation process prior to formation of an elongation-competent ribosome, the tilt of the 40S subunit is reversed, and the apex of domain II is released from its position on the head of the 40S subunit (Yamamoto et al., 2014). Remarkably, when levels of active eIF2 are reduced due to stress-induced phosphorylation, Met-tRNA_i_^Met^ can be recruited by eIF5B instead to the IRES/40S complexes (Pestova et al., 2008; Terenin et al., 2008). In both eIF2- and eIF5B-mediated pathways, eIF1A enhances 48S complex formation (de Breyne et al., 2008; Jaafar et al., 2016), whereas eIF1 inhibits the process and even induces dissociation of pre-assembled 48S ICs, but this inhibition can be alleviated by deletion of domain II (Pestova et al., 2008).

In addition to 40S subunits, HCV and related IRESs also bind to eIF3 via their apical IIIa and IIIb domains (Pestova et al., 1998b; Sizova et al., 1998; Ji et al., 2004; Hashem et al., 2013). Strikingly, in 40S/IRES/eIF3 complexes, HCV-like IRESs displace eIF3 from its ribosomal position (Hashem et al., 2013), usurping eIF3’s key ribosomal contacts involving eS1, eS26 and eS27 (des Georges et al., 2015). Moreover, the ribosome-binding surface of eIF3 is now involved in interaction with the IRES (Hashem et al., 2013). In *in vitro* reconstituted initiation reactions, eIF3 only modestly enhances 48S complex formation on HCV-like IRESs (Pestova et al., 1998b; Hashem et al., 2013), which led to the suggestion that *in vivo*, the role of the eIF3/IRES interaction is likely to relieve the competition between the IRES and eIF3 for a common ribosomal binding site, and to reduce formation of 43S PICs, thereby favoring translation of viral mRNAs (Hashem et al., 2013).

Cryo-EM studies have been indispensable in providing insights into the architecture and interactions of HCV and HCV-like IRES ribosomal complexes, as well as the mechanism of the IRES function, initially through low-resolution 40S/HCV IRES and 80S/HCV IRES structures (Spahn et al. 2001; Boehringer et al, 2005) and continuing with sub-nanometer resolution structures of 40S/eIF3/CSFV IRES and 80S/Met-tRNA_i_^Met^/eIF5B•GMPPNP/HCV IRES functional complexes (Hashem et al., 2013; Yamamoto et al., 2014), and the more recent near-atomic resolution reconstructions of 80S•HCV IRES complexes (Yamamoto et al., 2015; Quade et al., 2015; Yokoyama et al., 2019). However, despite these advances, the structures of 48S ICs assembled on the HCV IRES, as well as the transitions between different states in the initiation pathways and accompanying conformational changes have remained unknown. To fill these gaps, we present cryo-EM structures of HCV IRES ribosomal complexes up to 3.5 Å resolution that cover all major stages of IRES-mediated initiation pathways from IRES binding to the 40S subunit through eIF2-containing 48S ICs to the final eIF5B-containing 48S ICs immediately prior to the joining of the 60S subunit. Individually, these structures also provide detailed insights into the dynamic network of contacts between the IRES and the 40S subunit, highlight the role for IRES domain II, and importantly, include the first structure of eIF5B bound to the 40S subunit, prior to subunit joining.

## RESULTS AND DISCUSSION

### Overview of cryo-EM analysis of initiation complexes assembled on the wt and the ΔdII HCV IRES with eIF2 or eIF5B

To capture discrete states within either the eIF2- or eIF5B-containing IRES-mediated initiation pathways and to visualize the role of IRES domain II in these processes, initiation complexes were assembled *in vitro* by incubating the *wt* or the ΔdII mutant IRES (Figure 1A) with individual purified translation components. To follow the eIF2-mediated pathway, reaction mixtures were prepared containing the *wt* or the ΔdII IRES, 40S ribosomal subunits, eIF2, eIF3, eIF1A and Met-tRNA_i_^Met^, and to follow the eIF5B-mediated pathway, eIF2 was replaced by eIF5B, thus yielding four discrete sample types (Table S1). Cryo-EM grids of each complex were imaged at 300 kV producing high-contrast micrographs with easily identifiable 40S ribosomal particles (Figure S1A-E; Table S2). The images were processed using maximum-likelihood classification techniques implemented in Relion 3.1 (Scheres, 2012; 2016; Zivanov et al., 2018; 2019) yielding 18 structures containing different sets of components at resolutions as high as 3.5 Å (Figure S2; Table S3). Although some flexible regions had a poor local resolution (e.g., eS12 in the 40S head or IRES domain IIIb), most of the ribosome, all IRES-ribosome contacts, and all initiation factors present had resolutions, between 3-7 Å (Figures S2 and S3), that allowed modeling of all these components. None of the structures obtained contained eIF3. During initiation on HCV-like IRESs, eIF3 interacts with the apical region of IRES domain III rather than with the 40S subunit (Hashem et al., 2013). This interaction is sensitive to the process of grid preparation and is more stable when grids have thicker ice so that imaging complexes that contain eIF3 requires the intentional selection of regions with sufficiently thick ice (e.g., Hashem at al., 2013; Neupane et al., 2020); however, our study aimed to determine the details of ribosomal interactions with the IRES, initiation factors and Met-tRNAi^Met^ at high resolution, which relies on imaging in regions with thinner ice. Importantly, however, the absence of eIF3 does not affect data interpretation because all studied complexes can be assembled efficiently without eIF3 (Pestova et al., 1998b; 2008).

The structures obtained comprise the 40S/IRES_ΔdII_ binary complex in various conformational states (structures 1_ΔdII_-9_ΔdII_); the 40S/IRES_wt_ binary complex in a single conformational state (structure 10_wt_); the 40S/eIF1A/IRES_wt_ ternary complex (structure 11_wt_); 48S ICs containing eIF2, Met-tRNA_i_^Met^, eIF1A and the *wt* or the ΔdII IRES (structures 12_wt_ and 12_ΔdII_); 48S complexes containing the *wt* or the ΔdII IRES base-paired with the P-site Met-tRNA_i_^Met^ but lacking eIF2 and thus mimicking the stage after eIF2 dissociation following hydrolysis of GTP (structures 13_wt_ and 13_ΔdII_); the pre-48S IC containing eIF5B, eIF1A, the *wt* IRES and P-site Met-tRNA_i_^Met^ that is not base-paired with the initiation codon (structure 14 _wt_); and 48S ICs containing eIF5B, eIF1A, Met-tRNA_i_^Met^ and the *wt* or the ΔdII IRES (structures 15_wt_ and 15_ΔdII_) (Figure 1B). Thus, the structures obtained cover the entire initiation pathway, starting with initial binding of the IRES to the 40S subunit and finishing with the eIF5B-containing 48S complex prior to subunit joining, and also provide details of the dynamic interactions between the IRES and the 40S subunit (Figures 1C and S4; Table S4).

### Stepwise binding of the IRES to the 40S subunit

Binding of the IRES to the 40S subunit involves multiple contacts formed by several IRES domains (IIIa, IIIc, IIId, IIIe, S2) and ribosomal proteins eS1, eS27, eS28 as well as h26 in ES7 of 18S rRNA (e.g., Quade et al., 2015; Yamamoto et al., 2015). Deletion or mutation of these domains impairs binding of the IRES to the 40S subunit to different extents, reflecting their cumulative importance for IRES function (Kieft et al., 2001). Strikingly, classification of ribosomal complexes formed on the ΔdII IRES identified a small proportion of 40S/IRES binary complexes that showed conformational differences in the individual positions of IRES domains IIIa/IIIc/IIId/IIIe/IIIf (structures 1_ΔdII_-6_ΔdII_) compared to other complexes, in all of which these domains had similar positions (structures 7_ΔdII_-15_wt/ΔdII_) (Figure 1B). The local resolution of the IRES contacts with the 40S subunit in these maps was sufficient for model building of all components (Figures S2 and S3) and allowed detailed examination of the relative motion of the ΔdII IRES as it transitioned from a minimally associated state (structure 1_ΔdII_) to the canonically bound conformation in which domains IIIa, IIIc, IIId, and IIIe contact eS1, eS27 and h26 (structure 6_ΔdII_ and structures 7_ΔdII_-15_wt/ΔdII_).

By ordering these structures based on the number of IRES-40S subunit contacts and displacement from the canonically-bound IRES position we produced a putative sequence of binding events between the ΔdII IRES and the 40S subunit (Figure 2A; Table S4). Comparison between the least- and the most-bound states (structures 1_ΔdII_ and 6_ΔdII_) shows that the IRES domains undergo displacement of varying extents during IRES binding (Figure 2B). Across all structures, the most uniform regions of the IRES are the linked domains IIIa and IIIc (Figures 2A-B), which contact eS27 via nt. 163 and 233 as reported (Quade et al., 2015; Yamamoto et al., 2015). Given that these domains undergo minimal structural change regardless of the conformation of the rest of the IRES, the observed impairment of IRES activity by nucleotide substitutions (e.g., Tang et al., 1999) suggests that these interactions are critical for correct IRES function. Other IRES domains, however, are more dynamic. Thus, to transition from the least- to the most-bound states requires translation of domain IIId by 10.6 Å toward the intersubunit face and domains hIII_1_/IIIe/IIIf by 23.3 Å toward the mRNA exit channel, whereas domains IIIa/IIIc only move by 5.9 Å (Figure 2B).

**Figure 2.**
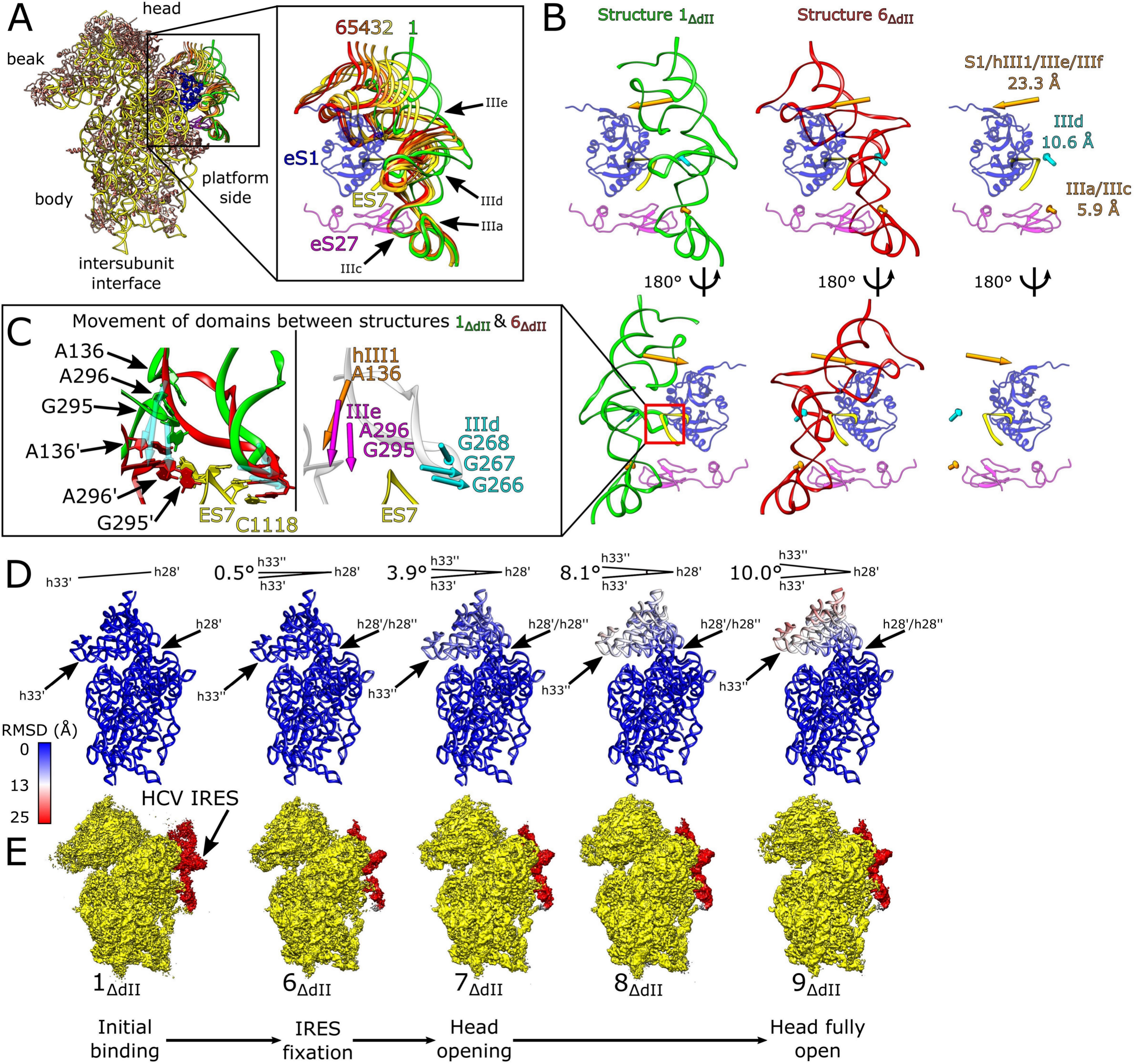
Initial events during binding of the HCV IRES to the 40S subunit. **(A)** IRES models from structures 1_ΔdII_-6_ΔdII_ aligned on the 40S subunit. IRES domains and ribosomal proteins eS1 (blue), eS27 (magenta) and 18S rRNA ES7 (yellow) are indicated (inset). **(B)** Comparison between minimally bound (structure 1_ΔdII_) and fully bound (structure 6_ΔdII_) IRES complexes showing the displacement of labeled IRES domains. **(C)** Formation of critical contacts between ES7 and the IRES requires reorganization of domains hIII_1_, IIIe, and IIId. The arrows indicate displacement of IRES nucleotides A_136_ (hIII_1_), G_295_/A_296_ (IIIe) and GGG_266-268_ (IIId) interacting with ES7 from the minimally bound (structure 1_ΔdII_) to the fully bound state (structure 6_ΔdII_) indicated with a prime (′). **(D)** RMSD (Å) of 18S rRNA for complexes labeled as in (E) compared to the minimally bound state (structure 1_ΔdII_), color-coded as in the inset key. The angle formed between helix 33 (h33ʹ) and helix 28 (h28ʹ) in structure 1_ΔdII_ and helix 33 (h33ʹʹ) in the labeled complex is also marked. **(E)** Segmented maps for labeled complexes showing 40S subunit (yellow) and IRES (red) organized in a putative sequence showing the minimally bound state (structure 1_ΔdII_), canonical IRES binding (structure 6_ΔdII_) and the induction of opening of the head of the 40S subunit (structure 7_ΔdII_-9_ΔdII_). See also Figure S4 and Table S4.

The fully bound IRES forms Watson-Crick base pairs between ES7 nt U_1114-1118_ and IRES domains IIId (GGG_266-268_), IIIe (A_296_), and hIII_1_ (A_136_) as well as a stacking interaction between domain IIIe (G_295_) and U_1115_ of ES7 (Quade et al., 2015; Yamamoto et al., 2015). This network of interactions also occurred in the later-stage complexes (structures 6_ΔdII_ and 7_ΔdII_-15_wt/ΔdI_) but they were not present during early-stage association of the IRES with the 40S subunit (structure 1_ΔdII_) (Figure 2C; summarized in Figure S4 and Table S4). Initially, the only contact between the IRES and ES7 is a transient and previously undescribed stacking pair between G_296_ in domain IIIe and U_1115_ of 18S rRNA. Structure 1_ΔdII_ also shows another transient, previously undescribed hydrogen bond between U_265_ in domain IIId and Lys199 in eS1, a residue that instead interacts with IIIe in the fully bound IRES (Quade et al., 2015; Yamamoto et al., 2015). These two contacts along with the domain IIIa/IIIc interactions with eS27 are the only bonds between the IRES and the 40S subunit in this complex (Table S4). In structure 2_ΔdII_, domain IIId begins to be repositioned, moving by 4.4 Å relative to structure 1_ΔdII_, and the transient domain IIIe-U_1115_ contacts are lost. Although the full complement of interactions with ES7 is not present, the Watson-Crick base-pairs between domain IIId (GG_266-267_) and CC_1117-1118_ and stacking interactions of G_295_ and A_296_ in domain IIIe with ES7 have formed. These Watson-Crick base pairs and the G_295_ stacking interaction are maintained in all subsequent (structures 2_ΔdII_-6_ΔdII_) and fully bound complexes (structures 7_ΔdII_-15_wt/ΔdI_), but the A_296_ stacking interaction with U_1115_ is only present for this and the following complex (structures 2_ΔdII_ and 3_ΔdII_). Another interaction that first appears in structure 2_ΔdII_ is the eS27 Glu75 hydrogen bond with domain IIId nts. 266-267. It is maintained in all subsequent structures except those that have domain II inserted into the E site of the 40S subunit (structures 10_wt_, 11_wt_ and 14_wt_; Table S4). In the following complex (structure 3_ΔdII_), domain IIId is located closer to the platform of the 40S subunit, moving by 2.8 Å; the S1/S2/hIII_1_/IIIe/IIIf region moves by 5.0 Å, whereas the position of the stable domain IIIa/IIIc changes by only 1.5 Å. This repositioning breaks none of the contacts formed in structure 2_ΔdII_ and allows the formation of a new base-pair between G_268_ in domain IIId and C_1116_ in ES7. The IRES domains continue to move closer to their canonical bound positions in structure 4_ΔdII_, which lacks the transient A_296_/U_1115_ interaction but maintains all other ribosomal contacts (Table S4). In structures 5_ΔdII_ and 6_ΔdII_, the final canonical interactions of hIII_1_ (A_136_) and domain IIIe (A_296_) with ES7 are present. The contact between A_136_ and U_1115_ is enabled because the base pairing between domain IIIe U_297_ and A_288_ in hIII_1_ that induces the flipping-out of A_136_ exists in all structures (1_ΔdII_-15_wt/ΔdII_) (Easton et al., 2009). Structures 5_ΔdII_ and 6_ΔdII_ are also the first complexes in which the IRES is in a position to form a hydrogen bond between Asn147 in eS1 and the phosphate backbone of GG_300-301_, an interaction that is maintained throughout all subsequent complexes (Table S4).

Taken together, this series of structures (1_ΔdII_-6_ΔdII_) indicates a likely sequence of binding events between the 40S subunit and the IRES from initial encounter to the canonically-bound conformation in which domain IIIa/IIIc, the first element of the IRES to bind to the 40S subunit, acts as a pivot to dock domain IIId onto ES7. These structures may represent transient states in binding of both the ΔdII and *wt* IRES, which we were able to capture in the former case because the altered kinetic landscape of the initiation pathway in the absence of domain II allowed them to accumulate and be observed.

An important corollary of IRES binding is the conformational changes that occur in the 40S subunit. Whereas complexes with an incompletely accommodated IRES (structures 1_ΔdII_-5_ΔdII_) contain 40S subunits in the analogous closed conformation, the ribosomal structures with the full complement of IRES/40S contacts (structure 6_ΔdII_-9_ΔdII_) show a striking difference between the position of the head, from the closed conformation in structure 6_ΔdII_ (matching the head position in structures 1_ΔdII_-5_ΔdII_) to the fully open state in structure 9_ΔdII_ (Figure 2D-E). Although structures 7_ΔdII_-9_ΔdII_ have a canonically bound IRES with domain III contacting eS1, eS27 and ES7 as in structure 6_ΔdII_, they show large-scale conformational changes to the head as the 40S subunit transitions from semi-closed (structure 7_ΔdII_) to fully open (structure 9_ΔdII_) states. Thus, structure 7_ΔdII_ opens by 3.9°, structure 8_ΔdII_ by 8.1°, and structure 9_ΔdII_ by 10.0° compared to the conformation of the 18S rRNA in structures 1_ΔdII_ -6_ΔdII_. These global changes to the position of the 40S head are reflected in changes in the P site as the distance between U_1248_ and C_1701_ increases from 7.3 Å, to 9.9 Å and finally to 11 Å in structures 7_ΔdII_, 8_ΔdII_ and 9_ΔdII_, respectively. Thus, this series of structures shows that even in the absence of domain II, establishment of the full complement of IRES/40S contacts results in the transition of the 40S subunit from the closed to the open state, which is required for accommodation of the initiation codon and surrounding regions in the mRNA-binding channel.

### Accommodation of the IRES in the mRNA-binding channel

In contrast to 40S/IRES binary complexes assembled on the ΔdII IRES, which showed remarkable differences in the position of the 40S subunit head, binary complexes assembled on the *wt* IRES yielded a uniform structure that was refined to 3.8 Å resolution from 119,320 particles (structure 10_wt_) (Figure 1B). Both the conformation of the 40S subunit and the structure of IRES domains IIIa-f in structure 10_wt_ are identical to those in the open state of the 40S/IRES_ΔdII_ binary complex (structure 9_ΔdII_) (Figures 3A-B). There is, however, additional density that corresponds to IRES domain II inserted into the 40S subunit E site, in a conformation that was observed in purified IRES/80S complexes (Quade et al., 2015; Yamamoto et al., 2015). Superposition of structure 10_wt_ and a closed-state 40S subunit shows that in the latter, steric clashes between uS7 and domain II would prevent insertion of this domain into the E site. Thus, domain II locks the 40S subunit into an open state, which is not similarly imposed in the case of the ΔdII IRES.

**Figure 3.**
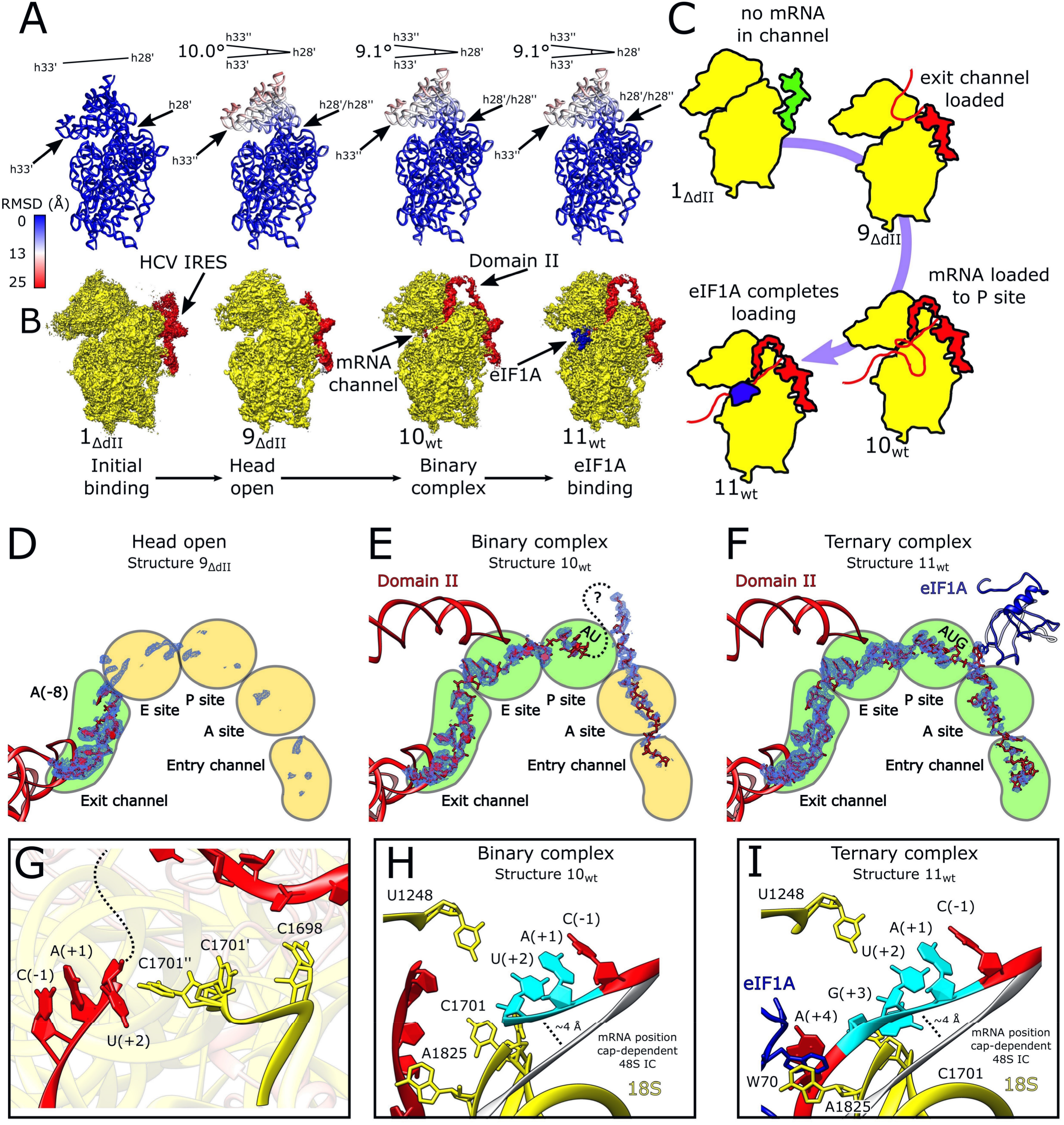
IRES domain II is required for loading mRNA into the mRNA channel. **(A)** RMSD (Å) of 18S rRNA for complexes labeled as in (B) compared to the minimally bound state (structure 1_ΔdII_), color-coded as in the inset key. The angle formed between helix 33 (h33ʹ) and helix 28 (h28ʹ) in structure 1_ΔdII_ and helix 33 (h33ʹʹ) in the labeled complex is also marked. **(B)** Segmented maps for labeled complexes showing 40S subunit (yellow), IRES (red), and eIF1A (blue) organized in a putative sequence showing the minimally bound state (structure 1_ΔdII_), fully opened head of the 40S subunit (structure 9_ΔdII_), binary complex (structure 10_wt_), and the eIF1A-containing ternary complex (structure 11_wt_). **(C)** Diagram of structures 1_ΔdII_, 9_ΔdII_, 10_wt_ and 11_wt_ showing the sequential loading of mRNA into the mRNA channel. **(D-F)** The mRNA channel spanning the entry and exit channels, A, P, and E sites viewed through the ribosome head towards the body for the labeled complexes. The IRES (red), Coulomb potential (blue mesh), and eIF1A (blue) are shown. **(G)** Position of key nucleotides in the P site in structure 10_wt_ showing interactions between incompletely loaded mRNA and C_1698_ resulting in C_1701_ sampling dual conformations near the (+2) position. **(H-I)** P site showing mRNA interactions for the (H) binary complex (structure 10_wt_) and the (I) eIF1A-containing ternary complex (structure 11_wt_). 18S rRNA (yellow), eIF1A (blue), IRES mRNA (red) with start codon (cyan) are all marked. The position of mRNA modelled in the 48S cap-dependent IC (PDB: 6ZMW) is shown in grey.

For the *wt* IRES, we also obtained the structure of the 40S•IRES•eIF1A ternary complex that was refined to 3.8 Å resolution from 204,320 particles (structure 11_wt_, Figure 1B). The conformation of the 40S subunit and the position of the IRES in it were identical to those in the 40S/IRES_wt_ binary complex (structure 10_wt_) (Figures 3A-B). The complex clearly showed density for eIF1A located between 18S rRNA helix (h) 44 and the ribosomal proteins eS30 and uS12, allowing us to model the OB domain and the C-terminal subdomain of eIF1A (residues 22-122). Although structure 11_wt_ lacks tRNA and the 40S subunit is in the open state, the position of eIF1A on the ribosome as well as its overall conformation are identical to those in the structures of eIF2-containing 48S complexes (Brito Querido et al., 2020; Simonetti et al., 2020).

As expected, binary complexes assembled on the ΔdII IRES and containing the 40S subunit in the closed conformation (1_ΔdII_-6_ΔdII_) do not have mRNA in the mRNA-binding channel. However, although structures 9_ΔdII_, 10_wt_ and 11_wt_ all have 40S subunits in the identical open conformation that is required for loading of mRNA into the channel, they differ strongly in the degree of ribosomal accommodation of the initiation codon and surrounding regions (Figure 3C). In the 40S/IRES_ΔdII_ binary complex (structure 9_ΔdII_), clear mRNA density was seen only in the exit portion of the channel up to the -8 position of mRNA (HCV nt 334), after which the mRNA became disordered (Figure 3D). In contrast, in the *wt* 40S/IRES binary complex (structure 10_wt_), mRNA nucleotides could be identified at the exit channel through the E site where it is stabilized by domain II, to AU_342-3_ located in the P site (Figure 3E). The (+3) nucleotide linked to U_343_ could not be identified due to disorder in the map. However, there is additional mRNA density in the mRNA channel ∼8 Å from the P-site G_344_, but the identity of these nucleotides does not correspond to those that immediately follow the start codon (A_345_ onwards) as this density extends 20Å out from the mRNA channel beyond the binding site of eIF1A, suggesting that after the P site, the mRNA is looped out (Figure 3E). Thus, the presence of domain II results in insertion of mRNA into the entire channel, but only the additional 9 nucleotides from the exit to the P site (i.e., from G_335_ to U_343_) can be reliably identified. Density corresponding to mRNA is present from the P site and entry channel but is likely a mixture of different registers of mRNA and so the sequence could not be determined (Figure 3E). Strikingly, the presence of eIF1A in ribosomal complexes resulted in accommodation of sequential mRNA along the entire mRNA binding cleft (Figure 3F). Examination of critical P-site nucleotides for all complexes shows that in each case C_1701_ of 18S rRNA is in a single conformation, except for the binary complex prepared with the *wt* IRES (structure 10_wt_) where it is present in two states as determined by examination of the Coulomb potential around that nucleotide (Figure 3G). In the conformation of the second state, C_1701_ contacts the upstream mRNA base U_343_, possibly contributing to stabilizing the mRNA when it has not undergone complete accommodation in the mRNA channel at the P site. The highly conserved C_1698_ (Prince et al., 1982) contacts downstream mRNA and may act as a sensor to stimulate C_1701_ adopting the second conformation in which it can stabilize the incompletely loaded mRNA (Figure 3G).

The position of the mRNA in structures 10_wt_ and 11_wt_, however, does not match its position in 48S complexes with an established codon-anticodon interaction (Brito Querido et al., 2020; Simonetti et al., 2020). Beginning from the (-1) position, the following (+1), and (+2) nucleotides are ∼4 Å above their position when tRNA is inserted (Figures 3H-I). However, the raised position of these nucleotides in the complex containing eIF1A nevertheless allows the (+4) adenine base to form the stacking triple with eIF1A Trp70 and 18S rRNA A_1825_ (Figure 3I), which is a key function of eIF1A (Battiste et al., 2000). These contacts are maintained up to formation of 48S complexes (Simonetti et al., 2020).

Taken together, the structures of binary complexes assembled on ΔdII and *wt* IRESs and the eIF1A-containing complex assembled on the *wt* IRES provide structural rationalization for the roles of domain II and eIF1A in sequentially loading the mRNA channel. Even without domain II, binding of the IRES induces conformational changes in the 40S subunit that are required for accommodation of mRNA in the binding channel. However, accommodation in this case is only partial, and the 40S subunit is not stably present in the open conformation, which is consistent, on one hand, with the ability of ΔdII IRES to function in initiation, but on the other hand, with lower initiation activity than the *wt* IRES (e.g., Reynolds et al., 1996). The presence of domain II results in accommodation of the initiation codon and the upstream region in the mRNA-binding channel, thereby enhancing the efficiency of initiation (Reynolds et al., 1996), whereas addition of eIF1A results in further accommodation of the mRNA in the entire mRNA-binding channel, accounting for its enhancement of initiation on the IRES (Jaafar et al., 2016).

### The structure of eIF2-containing 48S complexes assembled on the HCV IRES

The structures of eIF2-containing 48S ICs assembled on the *wt* and ΔdII IRES were refined from 46,904 and 103,813 particles to resolutions of 3.6 and 3.5 Å, respectively (structures 12_wt_ and 12_ΔdII_, Figures 1B and S1). Both *wt* and ΔdII IRES 48S ICs form identical complexes with respect to the 40S subunit’s closed conformation, the positions of Met-tRNA_i_^Met^, eIF2 and eIF1A, and the established P-site codon-anticodon base pairing (Figures 4A, and S5A-B). The IRES-containing 48S IC is also structurally identical to the canonical 48S complex formed by cap-dependent initiation (e.g., Simonetti et al., 2020) with respect to the global conformation of the 40S subunit as well as the placement of Met-tRNA_i_^Met^, eIF2α, and eIF1A. Thus, the E site-associated eIF2α contacts the highly conserved uS7/Asp194 directly and interacts with Met-tRNA_i_^Met^ via Thr103, Arg67 and His114 (Figure 4B). As with yeast 48S ICs (Hussain et al., 2014), eIF2α forms hydrogen bonds with the (-3) adenosine via Arg55, a contact that enhances codon selection in the scanning mode of initiation, presumably by stabilizing the arrested 48S IC (Pisarev et al., 2006; Thakur et al., 2020). Contacts between eIF2α Arg57 and the tRNA acceptor stem loop (ASL) and the 18S rRNA that occur in canonical 48S ICs are also present (Simonetti et al., 2020) (Figure 4B). The position of eIF2α is identical in *wt* and ΔdII IRES 48S ICs (Figure S5E). The unsharpened map also contained density corresponding to eIF2γ (Figure S5D) identical to that seen in cap-dependent 48S initiation complexes (Simonetti et al., 2020), but it had a low local resolution at the acceptor end of Met-tRNA_i_^Met^ and was not modelled after map sharpening.

**Figure 4.**
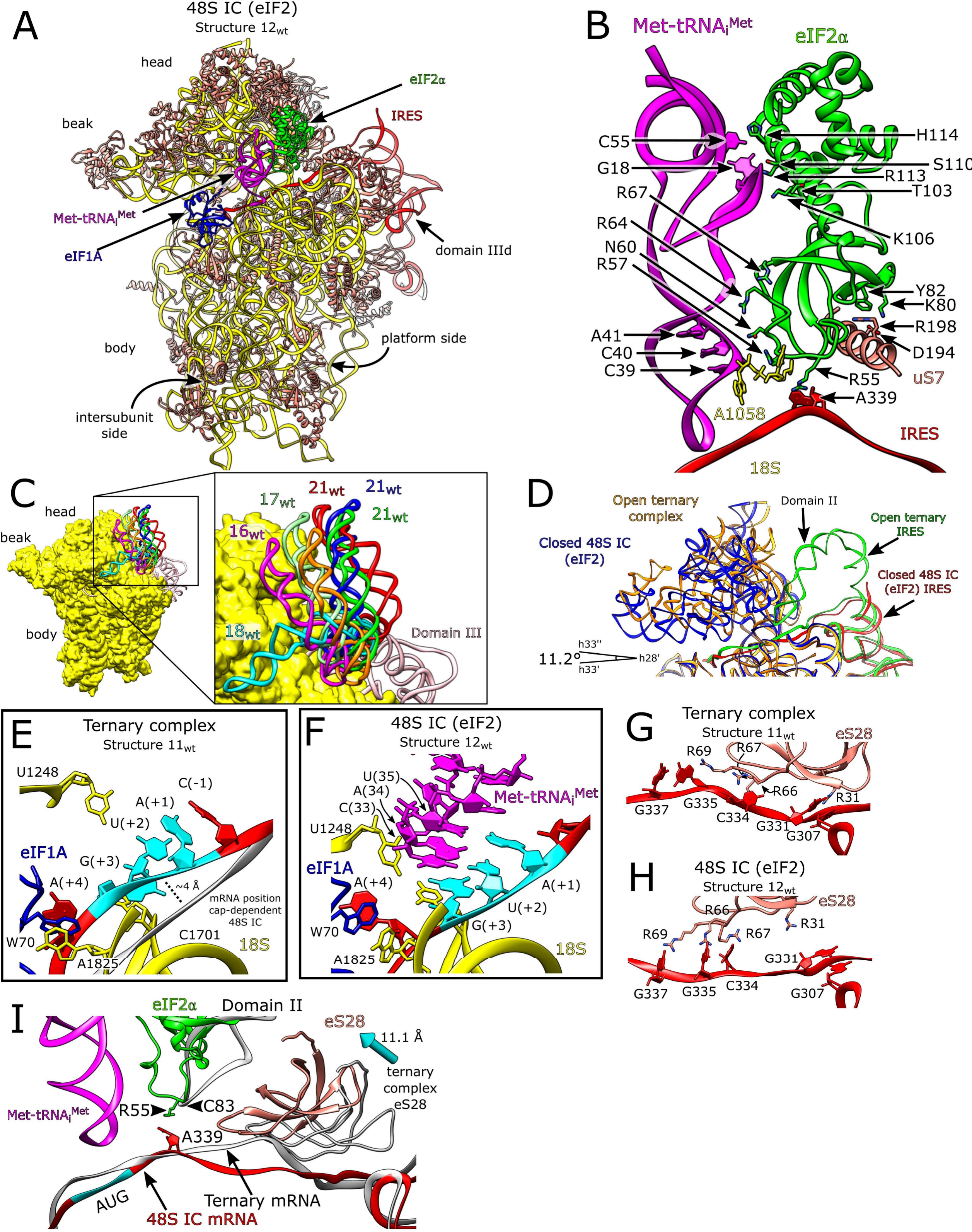
The HCV IRES•eIF2-containing 48S initiation complex. **(A)** Overview of the *wt* IRES eIF2-containing 48S IC (structure 12_wt_). **(B)** Contacts between eIF2α and HCV IRES, Met-tRNAi^Met^, and 40S subunit. **(C)** IRES domain II occupies multiple positions in the eIF2-containing 48S IC. **(D)** Position of 18S rRNA in the open ternary complex (structure 11_wt_) and eIF2-containing 48S IC (structure 12_wt_). **(E-F)** The P site in the (E) ternary complex (structure 11_wt_) and the (F) eIF2-containing 48S IC (structure 12_wt_) showing 18S rRNA (yellow), Met-tRNAi^Met^ (magenta), eIF1A (blue), and IRES mRNA (red) with the start codon (cyan) marked. **(E)** The position of mRNA in the cap-dependent 48S IC is marked (grey). **(G-H)** Contacts between eS28 (salmon) and the IRES (red) in (G) the ternary complex (structure 11_wt_) and (H) the eIF2-containing 48S IC (structure 12_wt_). **(I)** Global conformation of mRNA in the ternary complex (structure 11_wt_) and the eIF2-containing 48S IC (structure 12_wt_). The arrow indicates the extent of movement of eS28 between the binary complex (structures 11_wt_) and the eIF2-containing 48S IC complex (structure 12_wt_). See also Figure S5 and Table S5.

The position of domain II differs substantially between the 48S IC assembled on the *wt* IRES (structure 12_wt_) and the corresponding 40S/IRES binary and 40S/eIF1A/IRES ternary complexes (structures 10_wt_ and 11_wt_). In both the binary and ternary complexes, it is inserted into the E site, a position that is incompatible with the binding of eIF2. Thus, in the 48S IC, domain II is oriented away from the subunit interface, towards the solvent side of the 40S subunit, and shows an attenuation of density so that a model of domain II is not fully enclosed by the map. Focused classification of this region revealed that domain II is flexible and occupies multiple conformations oriented away from the E site (Figure 4C; Figure S5F; Table S5).

Compared with the binary or ternary complexes (structures 10_wt_ and 11_wt_), the head of 48S ICs formed on both *wt* and ΔdII IRESs is in a closed position, having moved by 11.2° relative to the open states (Figure 4D), and is in an even more closed conformation than in closed binary complexes (structures 1_ΔdII_-6_ΔdII_), in which the position of the head differed from that in structures 10_wt_ and 11_wt_ by ∼9.0° (Figure 3A).

Comparison between the ternary complex (structure 11_wt_) and the 48S IC (structure 12_wt/ΔdII_) shows the effect that incorporation of tRNA into the P site and rearrangement of 18S rRNA into the closed conformation has on the position of mRNA in the mRNA-binding channel. The raised position of P-site mRNA in the ternary complex (Figure 4E) cannot be maintained as this would cause a clash with the P-site tRNA in the 48S IC (Figure 4F). Thus, upon 40S subunit closure and incorporation of P-site tRNA, the mRNA repositions ∼4 Å deeper into the mRNA channel and the A(+4) base flips out to maintain the stacking triple with eIF1A Trp70 and 18S rRNA A_1825_ that is seen in the ternary complex (Figures 4E-F). The mRNA is also shifted upstream due to the movement of eS28 in the ribosome head (Figure 4G). A network of hydrogen bonds in the ternary complex (structure 11_wt_) between multiple arginines in eS28 and the mRNA is reformed by the displacement of eS28 in the closed complex (structure 12_wt/ΔdII_). Arg69 contacts the guanine at (-7) position in the ternary complex, but the shift of the 18S rRNA to the closed position causes this leading contact to be broken and reformed with G(-5). Other contacts are similarly reorganized, and the Arg31 contact with the stacked G_331_-G_307_ pair is lost completely (Figure 4G-H). This reorganization of contacts causes the mRNA to shift by one base pair at the P site and allows contact with the tRNA anticodon arm once tRNA is inserted into the P site. Interestingly, eIF2α Arg55 contacting the mRNA at A(-3) is in the same position as C_83_ in IRES domain II in the 40S/eIF1A/IRES ternary complex, which could indicate that this location in the E site is important for stabilizing the mRNA regardless of the conformation of the 40S subunit and differences in position of mRNA at other locations along the mRNA-binding channel (Figure 4I). Interestingly, we also obtained 40S ribosomal complexes containing platform-bound *wt* or ΔdII IRES and P-site Met-tRNA_i_^Met^ but lacking all initiation factors (structures 13_wt_ and 13_ΔdII_) (Figure 1C). These closed complexes, refined from 15,906 and 15,598 particles to 4.6 and 4.4 Å, respectively, clearly showed density for codon-anticodon base pairing (Figure S3G; Table S3). It is unclear whether initiation factors dissociated due to slow eIF5-independent hydrolysis of eIF2-bound GTP (Unbehaun et al., 2004), or through denaturation at the air-water interface and/or due to the shear forces associated with blotting (Glaeser, 2021; d’Imprima et al., 2019). In any case, this complex likely mimics an intermediate state immediately after eIF2 dissociation and prior to the binding of eIF5B.

### The structure of eIF5B-containing 48S complexes assembled on the IRES

Classification of eIF5B-containing ribosomal complexes yielded two structures formed on the *wt* IRES (structures 14_wt_ and 15_wt_) and one structure (15_ΔdII_) assembled on the ΔdII IRES (Figures 1B and S1). Structures 14_wt_, 15_wt_ and 15_ΔdII_ were refined to 3.8 Å, 3.7 Å and 3.7 Å resolution from 60,578, 133,782 and 61,648 particles, respectively (Figures 5A-B and S6A). All structures contain eIF5B, eIF1A and the P-site Met-tRNA_i_^Met^. However, whereas structures 15_wt_ and 15_ΔdII_ showed density for the base-paired codon-anticodon and were accordingly classified as 48S ICs, the P-site Met-tRNA_i_^Met^ was not base-paired with the initiation codon in structure 14_wt_, which was therefore designated as a pre-48S IC (Figure S3G). Similar to eIF2-containing 48S ICs assembled on the *wt* IRES, in eIF5B-containing ICs assembled on the *wt* IRES (structure 15_wt_) domain II is oriented away from the subunit interface, toward the solvent side of the 40S subunit, and shows an attenuation of density that can be resolved by focused classification into multiple states (Figure 1B and S6B; Table S5). In contrast to structures 15_wt/ΔdII_ that are in an identical closed conformation, the 40S subunit in structure 14_w_ is in an open conformation with domain II inserted into the E site (Figure S6C). In all structures, eIF1A occupies its usual position over h44 and eS30 and uS12, whereas eIF5B resides on the intersubunit face of the 40S subunit as in 80S ribosomal complexes (Yamamoto et al., 2014; Huang and Fernandez, 2020; Wang et al., 2020).

**Figure 5.**
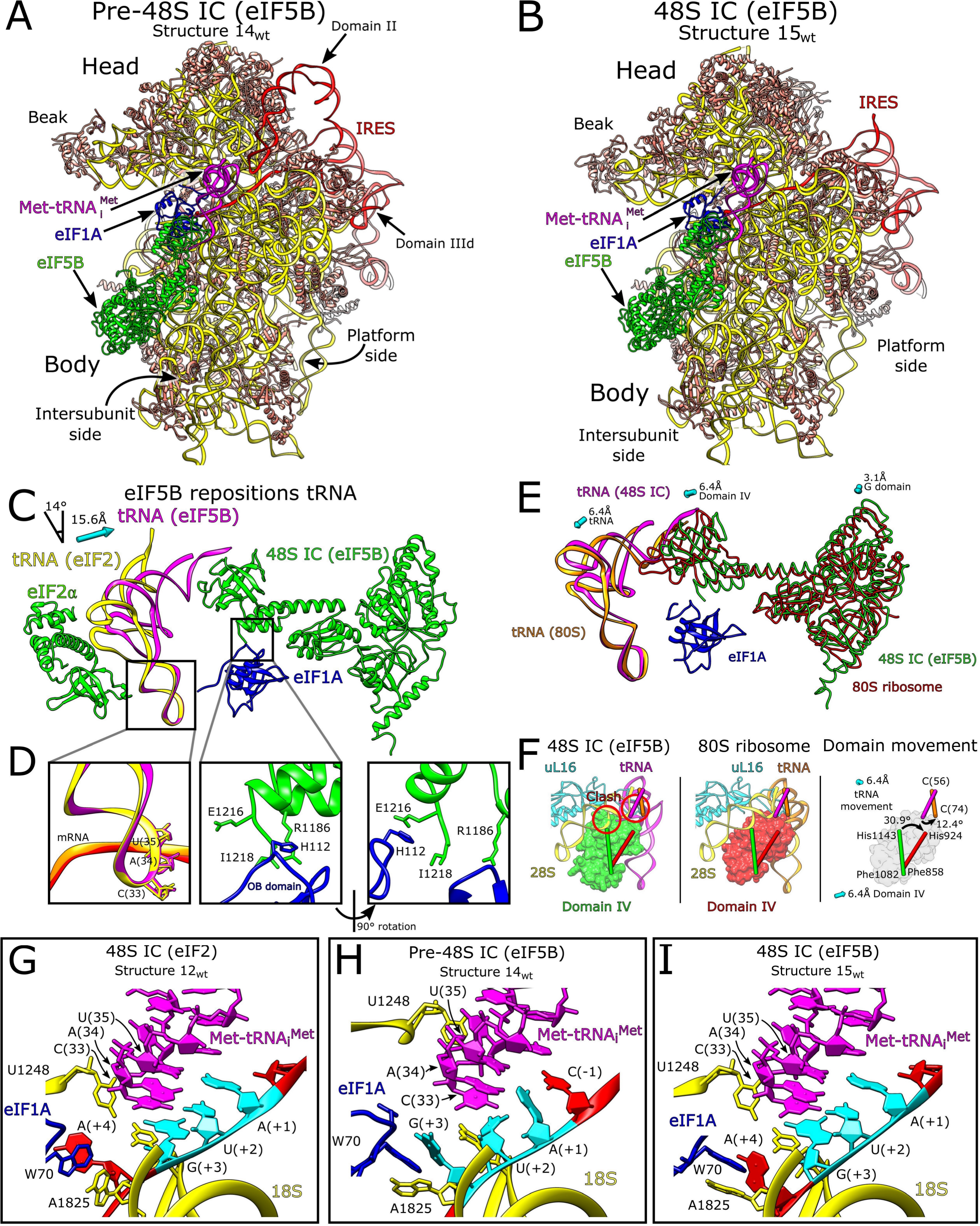
The HCV IRES•eIF5B-containing 48S initiation complex. **(A)** Overview of the *wt* IRES eIF5B-containing pre-48S IC (structure 14_wt_). **(B)** Overview of the *wt* IRES eIF5B-containing 48S IC (structure 15_wt_). **(C)** Changes to the position of P site tRNA depending on the presence of either eIF2 or eIF5B. Upon binding of eIF5B, the T-arm, D-arm and acceptor stem of Met-tRNAi^Met^ move by 15.6 Å and 14° relative to their positions in the eIF2-containing 48S complex. **(D)** Inset showing the conformation of tRNA acceptor stem loop, and contacts between eIF5B domain IV and eIF1A. The conformation of the ASL is unchanged between the pre-48S IC (eIF5B), and the 48S IC (eIF2 or eIF5B). **(E)** Position of eIF5B and tRNA in the 48S and pre-elongation 80S ribosome complexes. eIF5B undergoes relatively little movement between the 48S (green) and 80S stages (red) in the II, G, and III domains, but domain IV translates and rotates (see F) causing movement in the tRNA between the two complexes (magenta and orange respectively). Arrows show displacement for labeled domains or components between the eIF5B-containing 48S IC and pre-elongation 80S ribosome. **(F)** The position of eIF5B domain IV in the 48S IC (green) would clash with 28S rRNA H89 and uL16 in the 60S subunit. Upon binding of the 60S subunit, eIF5B domain IV (red) translates by 6.4Å towards to the platform side of the 40S subunit and rotates by 30.9° causing the tRNA to rotate by 12.4° and translate by 6.4Å towards the head. **(G-I)** P site for (G) eIF2-containing 48S IC (structure 12_wt_), (H) eIF5B-containing pre-48S IC (structure 14_wt_), and (I) eIF5B-containing 48S IC (structure 15_wt_) showing 18S rRNA (yellow), Met-tRNA_iMet_ (magenta), eIF1A (blue), and IRES mRNA (red) with the start codon (cyan) marked. See also Figures S6-S7 and Tables S5-S6.

The structures show clear density for eIF5B residues 592-1218, corresponding to all major domains. The G domain, and domains II and III form the central domains of eIF5B that connect via domain III and helix 12 to the tRNA acceptor stem binding domain IV. (Figure 5C). This is the first high-resolution structure of mammalian eIF5B and also the first structure of the 40S/eIF5B subunit complex prior to subunit joining. Although we used full-length native eIF5B, extensive 3D classification and processing did not reveal any structure corresponding to the N-terminal region, suggesting that the structure in this region is highly disordered, an observation that is supported by the structure of full-length eIF5B from a range of species predicted using AlphaFold (Jumper et al., 2021). Consistent with the requirement of the 60S subunit for induction of eIF5B’s GTPase activity (Pestova et al., 2000), all complexes contained eIF5B-bound GTP (Figure S6E). Previous structures of mammalian eIF5B determined at ∼9 Å resolution used GMPPNP and showed the GDP-bound conformation of switch 1 (Yamamoto et al., 2014) (Figure S7A). In contrast, all our complexes showed switch 1, switch 2, and the β9-β10 loop in domain II in the GTP-bound conformation (Kuhle and Ficner, 2014). The conformation of this conserved GTP-binding region of mammalian eIF5B is identical to that in fungal eIF5B (Kuhle and Ficner, 2014; Wang et al., 2020) (Figure S7B-D).

We identified multiple contacts between eIF5B domains II and III and 18S rRNA h5, h14, and h15, as well as an interaction between eIF5B domain II and uS12 (Table S6; Figure S6D), and noted that the β13-β14 loop in our complexes could interfere with the transition of switch 1 from GTP-bound to GDP-bound conformations (Kuhle and Ficner, 2014) (Figure S6F). On the yeast 80S ribosome, the β13-β14 loop of eIF5B contacts A_415_ in 18S rRNA h14 and is positioned away from the path that switch 1 might take as it changes to the GDP-bound conformation (Wang et al., 2020). Although there is no contact between the equivalent nucleotide (A_464_) and β13-β14 in our complexes, h14 is accessible. These observations suggest that the 60S subunit might stimulate the interaction between A_415_ and the β13-β14 loop to reposition this loop away from switch 1 so that a transition from GTP- to GDP-bound conformation. In both pre-48S ICs and 48S ICs, the position of domain IV also allows it to contact eIF1A via interactions between His112 in the helical subdomain of eIF1A (Battiste et al., 2000) and the extreme C-terminal region of eIF5B, as well as between the eIF1A L23 beta-turn (near Gly54) and Arg1186 in domain IV of eIF5B (Figure 5D).

These interactions are distinct from the previously reported binding of eIF1A’s extreme C-terminal DDIDI sequence to the h12/h13 loop in eIF5B domain IV (Marintchev et al., 2003; Zheng et al., 2014) and of the eIF1A L45 loop to eIF5B domain III (Nag et al., 2016).

Comparison of the position of the P-site Met-tRNA_i_^Met^ in the eIF2-containing 48S IC (Simonetti et al., 2020) and in the 80S ribosome (Yamamoto et al., 2014; Wang et al., 2020) shows when the large subunit is present, tRNA rotates 14° towards the 40S subunit body and the T-loop moves by ∼ 15 Å to allow placement of the acceptor stem into the P site of the 60S subunit. We therefore examined our maps (structures 12_wt/ΔdII_-15_wt/ΔdII_) to determine how the orientation of the P-site tRNA differs in 40S ribosomal complexes depending on the presence of eIF2 or eIF5B. Compared to eIF2-containing 48S ICs, in eIF5B-containing complexes, Met-tRNA_i_^Met^ rotates by ∼14° and moves by 15 Å from the head of the 40S subunit to a position that matches the orientation seen in 80S structures (Wang et al., 2020) (Figure 5C). This repositioning of tRNA was observed for all structures that contained eIF5B, regardless of whether the 40S subunit was in the open (pre-48S IC) or closed (48S IC) conformation (Figure 5C), indicating that eIF5B re-orients Met-tRNA_i_^Met^ on the 40S subunit at the 48S PIC stage prior to subunit joining. Interestingly, although the overall position of eIF5B-Met-tRNA_i_^Met^ in eIF5B-containing 48S ICs and in 80S ribosomes is similar, upon binding of a 60S subunit, domain IV of eIF5B undergoes a 33° rotation towards the platform side of the 40S subunit as well as a translation by 6.4 Å parallel to the mRNA channel towards the platform of the 40S subunit, which results in repositioning of the tRNA acceptor stem by 6.4 Å toward the ribosome head (i.e., toward ribosomal protein eS25) without changing the position of the ASL (Figure 5E-F). The repositioning of eIF5B domain IV occurs despite an only minor movement of helix 12 (Figure 5F; Figure S6G). Such repositioning of domain IV and the tRNA acceptor stem upon binding of a 60S subunit would avert steric clashes of domain IV with H84 of 28S rRNA and between uL16 and the tRNA acceptor stem (∼A74) (Figure 5F). Similar repositioning of IF2 domain C2 (equivalent to eIF5B domain IV) to avoid analogous steric clashes was observed in bacteria upon joining of 50S subunits to 30S ICs (Hussain et al., 2016). Adjustment of the orientation of initiator tRNA prior to subunit joining is a critical step in initiation, and the mechanism by which it is mediated, by rotation and translation of domain IV of the universally conserved initiation factor IF2/eIF5B, likely appeared early in evolution.

Despite the differences in the orientation of the acceptor arm of the P-site tRNA in ribosomal complexes containing eIF2 and eIF5B, as well as differences in the conformation of the 40S subunit in eIF2/eIF5B-containing 48S ICs and eIF5B-containing pre-48S ICs, the anticodon loop in all these complexes is identically positioned (Figure 5D). eIF5B- and eIF2-containing 48S ICs assembled on the IRES are also in the same closed conformation (Figures S6C), identical to that in 48S ICs assembled on canonical cap-dependent mRNA (Simonetti et al., 2020). Thus, in eIF5B- and eIF2-containing 48S ICs, 18S rRNA nucleotides C_1701_ and U_1248_ are separated by ∼3.5 Å, and the contacts of the P-gate nucleotides G_1639_ and A_1640_ in the 18S rRNA and Arg146 in uS9 with the tRNA anticodon arm on the opposite side to the anticodon are also present in both 48S ICs.

In contrast, the 40S subunit in the eIF5B-containing pre-48S IC is in the open conformation, in which the separation between C_1701_ and U_1248_ is increased to ∼11 Å, and there are no contacts between the tRNA and the P gate or uS9 (Figure 5H). As with the 40S/eIF1A/IRES ternary complex (structure 11_wt_), the open configuration of the ribosome head present in the pre-48S IC creates a network of hydrogen bonds between the mRNA and eS28 (Figure S7E) that causes the mRNA to be shifted by one nucleotide relative to the closed configuration (Figure S3G). This results in the final nucleotide of the start codon (G_344_) being out of place in the P site and not participating in any contacts with the tRNA anticodon. Instead, G_344_ is available to form a stacking triple with eIF1A Trp70 and A_1825_ of 18S rRNA (Figure 5H). In the closed conformation of 48S ICs, the hydrogen bonds with eS28 have been partially broken (Figure S7F) and so all three nucleotides of the start codon are accommodated in the P site, and the A(+4) nucleotide is available to form a stacking triple with eIF1A Trp70 and A_1825_ of 18S rRNA (Figure 5F and 5I).

### Comparison of eIF5B-containing pre-48S initiation complexes with eIF2-containing scanning 43S complexes

In contrast to initiation on the HCV IRES, eIF5B cannot substitute for eIF2 in recruiting Met-tRNA_i_^Met^ to the 40S subunit in the canonical scanning mechanism. Canonical initiation also requires eIF1, which binds to the 40S subunit below the mRNA channel at the P site between h24 and the region connecting h44 to h45 and, in cooperation with eIF1A, induces the open conformation of the 40S subunit (Passmore et al., 2007; Llácer et al., 2015), thereby promoting ribosomal attachment to mRNA, scanning and initiation codon selection (e.g., Pestova and Kolupaeva, 2002). We therefore compared the conformation of the 40S subunit and the positions of mRNA and Met-tRNA_i_^Met^ in eIF5B-containing pre-48S complexes formed on the *wt* IRES (structure 14_wt_) and in canonical scanning eIF2-containing 43S complexes that also contained eIF1 (Brito Querido et al., 2020).

Even though the individual mRNA bases could not be identified in the scanning, eIF2-containing 43S complex because the P site nucleotides were heterogenous (Brito Querido et al., 2020), the positions of mRNA in this complex and in the eIF5B-containing pre-48S complex are very similar (Figure 6A). Although codon-anticodon base-pairing is not established in either complex, the anticodon loops of Met-tRNA_i_^Met^ in both are poised to contact mRNA and base-pair with an initiation codon, which is in contrast to the P_OUT_ position of tRNA in fully open mRNA-free 43S complexes (Kratzat et al., 2021), in which the anticodon loop is positioned too far away to be able to interact with mRNA if it had bound in the P site (Figure S7G). However, whereas the conformations of the anticodon in the P site in both complexes are similar, in the eIF5B-containing complex, the T-arm and the acceptor stem are shifted toward the body of the 40S subunit by bending of the anticodon loop region (Figure 6B). Consequently, the contacts between the ASL of Met-tRNA_i_^Met^ and 18S rRNA nucleotides GA_1639-40_ and the N-terminal region of uS9 in the head of the 40S subunit, which are present in all eIF2-containing complexes, including mRNA-free and scanning 43S complexes and 48S ICs (e.g., Hussain et al., 2014; Brito Querido et al., 2020; Simonetti et al., 2020), do not exist in the eIF5B-containing pre-48S ICs and only form after codon-anticodon base-pairing (Figure 6B). Thus, interaction with eIF2 allows tRNA to maintain the contacts with the head of the 40S subunit in all conformations, from fully open to fully closed, and the position of the anticodon loop that allows it to inspect mRNA in the scanning 43S complexes is determined by the rotation of the head that is in the intermediate conformation compared to fully open and fully closed states. In contrast, when interacting with eIF5B, tRNA establishes these contacts only in the closed position of the head, whereas in the open pre-48S ICs tRNA is stabilized by contacts between eIF5B domain IV and the acceptor stem, tRNA anticodon U_35_ and mRNA A(+1), and a single 18S rRNA nucleotide (C_1701_) that partially stacks with C_33_ (instead of the stacking of both U_1248_/C_1701_ as in the closed case) (Figure 6B). Thus, whereas some aspects of eIF5B-containing pre-48S complexes are analogous to those of eIF2-containing scanning 43S complexes, the overall orientation, and the specific interactions of tRNA in them differ.

**Figure 6.**
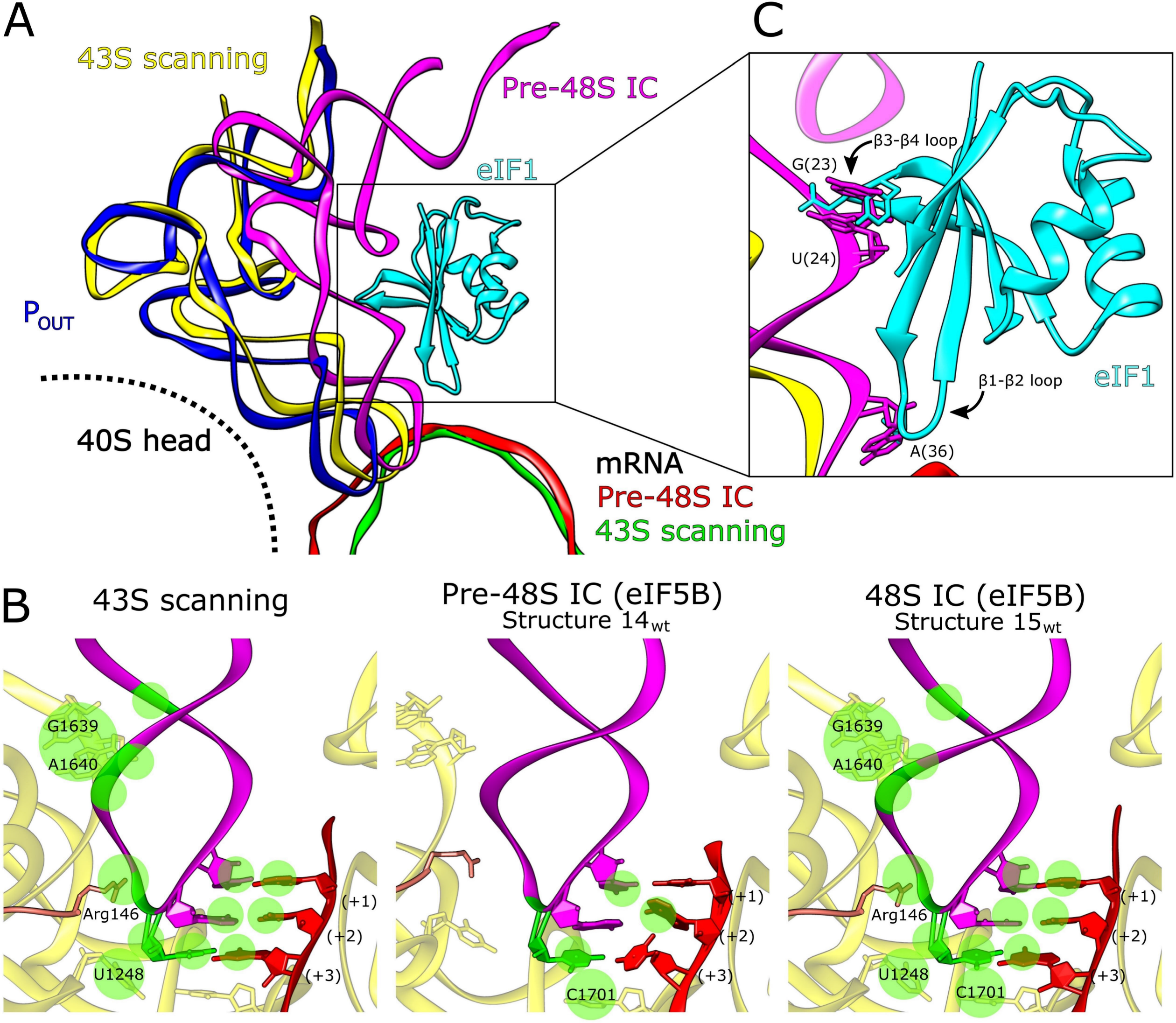
Position of mRNA in the ribosomal P-site and codon-anticodon recognition. **(A)** Position of tRNA in the P site for the P_OUT_ complex (PDB: 7A09) (blue), 43S scanning complex (PDB: 6ZMW) (yellow), and the pre-48S IC (structure 14_wt_) (magenta). mRNA from the 43S scanning complex (green) and pre-48S IC (red) is shown. **(B)** Contacts between tRNA, the initiation codon and the P site for the labeled complexes. **(C)** The β1-β2 and β3-β4 loops of eIF1 bound as in the 43S scanning complex (PDB: 6ZMW) would clash with the pre-48S IC position of Met-tRNAi^Met^ . See also Figure S7.

In eIF5B pre-48S complexes, the open conformation of the 40S subunit is supported by the insertion of IRES domain II into the E site, whereas in eIF2-containing complexes, the conformation of the 40S subunit is determined by the binding of eIF1. We therefore analyzed whether binding of eIF1 would be compatible with the structure of eIF5B-containing pre-48S ICs. The position of eIF1 placed into such complexes suggests that it would clash with tRNA. Thus, the repositioning of tRNA in eIF5B-containing complexes causes the AAC_38-40_ nucleotides of Met-tRNA_i_^Met^ to move toward the 40S subunit body by ∼3.0 Å, so that the binding of eIF1 as in the 43S scanning complex would create a clash between A_36_ of Met-tRNA_i_^Met^ and the β1-β2 loop (Figure 6C). Accommodation of eIF1 would require either reorganization of this loop or displacement of the P-site tRNA. Examination of human (Fletcher et al., 1999) and yeast (Reibarkh et al., 2008) solution NMR structures of eIF1 did not identify any conformations of the β1-β2 loop that would allow a clash with the anticodon stem of tRNA to be avoided in the eIF5B-containing pre-48S and 48S ICs. Moreover, the eIF1 β3-β4 loop would also clash with tRNA nucleotides GU_23-24_, and a clash between Phe113 and tRNA nucleotide G_25_ is also possible (Figure 6C). These observations suggest that even if eIF5B were able to bind Met-tRNA_i_^Met^ with high affinity and recruit it to the 40S subunit efficiently, the structure of the resulting complexes would not be compatible with binding of eIF1 and hence, with the scanning mechanism of initiation. On the other hand, the stabilizing interaction of the acceptor arm of Met-tRNA_i_^Met^ with domain IV of eIF5B in the closed 48S complexes following dissociation of eIF2•GDP and eIF1 would lock the complex preventing leaky scanning from occurring.

The incompatibility of Met-tRNAi^Met^ and eIF1 on eIF5B-containing pre-48S complexes likely explains why eIF1 disrupts 48S complexes prepared using *wt* but not the ΔdII variant of the HCV-like CSFV IRES (Pestova et al., 2008). HCV domain II has the propensity to insert into the E site (Quade et al., 2015; Yamamoto et al., 2015), locking the ribosome into the open conformation, and the structurally related CSFV domain II likely behaves similarly. As outlined above, the tRNA consequently loses contacts with uS9, with GA_1639-40_ as well as with U_1248_. If eIF1 binds to this complex, then the insertion of its β1-β2 loop into the mRNA channel creates steric hindrance between mRNA and tRNA in the P site, dislodging the tRNA.

### CONCLUDING REMARKS

Here we present the most comprehensive structural overview of the HCV IRES-mediated initiation pathway to date (Figure 7). The IRES initially binds to the 40S subunit through domains IIIa/IIIc and then pivots onto its platform side where it establishes the complete set of contacts (structures 1_ΔdII_-6_ΔdII_). Once the canonical set of contacts are made, this induces the head of the ribosome to open (structure 8_ΔdII_-9_ΔdII_). Although head opening can occur in the absence of IRES domain II, such complexes are nevertheless characterized by remarkable heterogeneity in the position of the 40S subunit head. In contrast, 40S/IRES binary complexes assembled on the *wt* IRES yield a uniform structure, in which the 40S subunit is in the open conformation, and domain II is inserted into the E site (structure 10_wt_). Importantly, in the absence of domain II, mRNA density was clearly seen only in the exit portion of the channel up to the -8 position of mRNA (structure 9_ΔdII_), whereas in the *wt* 40S/IRES binary complex, mRNA nucleotides could be identified at the exit channel through the E site where it is stabilized by domain II, to AU_342-3_ located in the P site (structure 10_wt_), and eIF1A induces further accommodation of the mRNA in the entire mRNA-binding channel (structure 11_wt_). Thus, these complexes provide structural insights into the functions of multiple IRES domains, including IIIa/IIIc in establishing the initial ribosome contacts, IIId in fixing the IRES to the 40S subunit and inducing ribosomal head opening, and II in imposing the open conformation and promoting fixation of mRNA in and upstream of the P site. Our analysis also revealed the role of eIF1A in completing mRNA accommodation.

**Figure 7.**
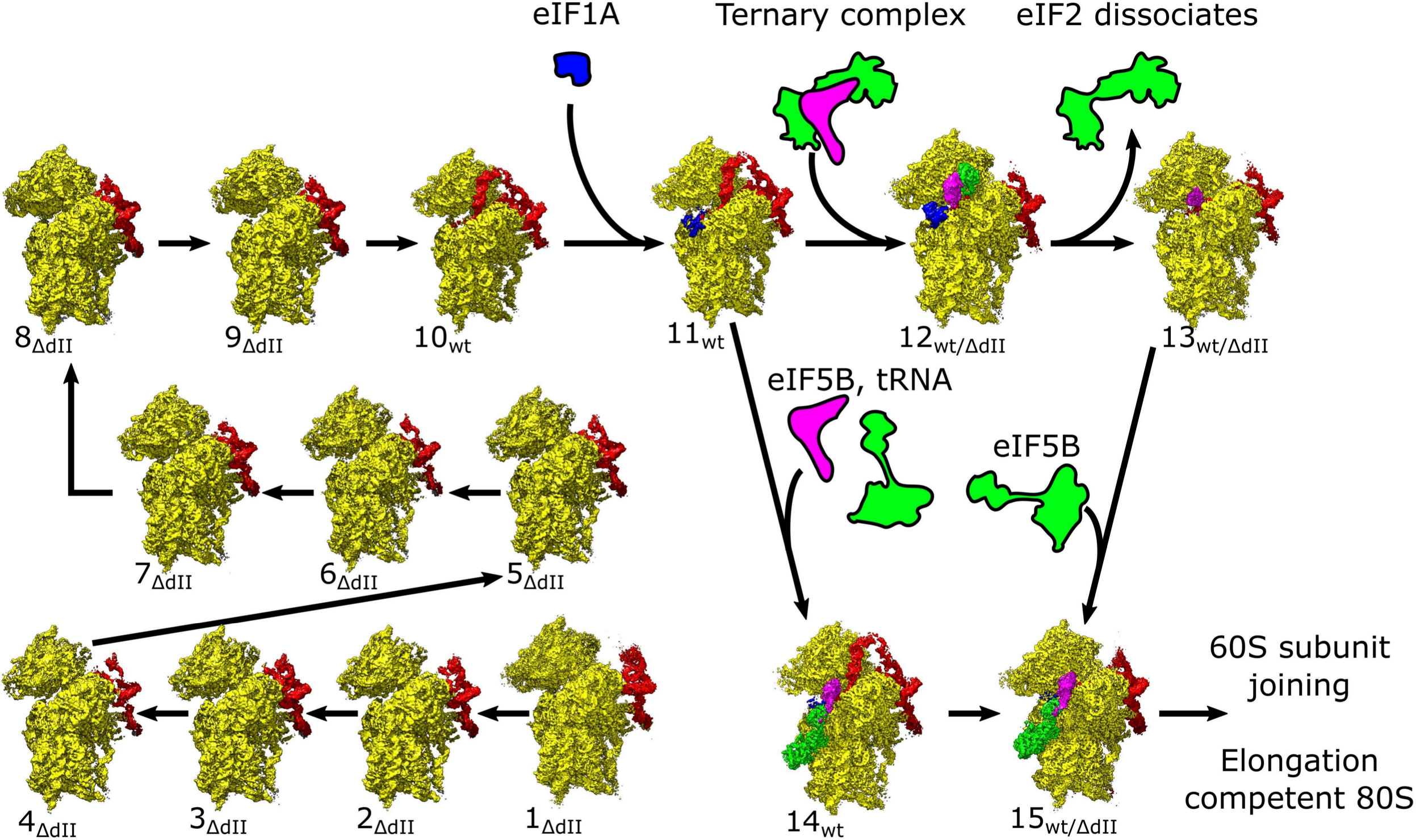
The HCV IRES-mediated initiation pathway. Ribosomal complexes organized in a putative IRES-mediated initiation pathway. Maps are segmented showing the 40S subunit (yellow), IRES (red), eIF1A (blue), Met-tRNAi^Met^ (magenta), and eIF2 or eIF5B (green).

Once the mRNA is loaded, initiation can proceed either along the canonical initiation pathway, in which eIF2 promotes attachment of Met-tRNA_iMet_ to form the 48S IC (structure 12_wt/ΔdII_) and then dissociates after GTP hydrolysis (likely structure 13_wt/ΔdII_) followed by binding of eIF5B (structure 15_wt/ΔdII_), or by a shortcut route, in which Met-tRNA_iMet_ is loaded directly by eIF5B, first forming open pre-48S ICs without established codon-anticodon interaction (structure 14_wt_) and then locking upon codon-anticodon base-pairing (structure 15_wt_). The eIF2-containing 48S ICs assembled on the IRES are structurally identical to canonical 48S ICs with respect to the conformation of the 40S subunit and the positions of Met-tRNAi^Met^, eIF2 and eIF1A. The position of domain II of the IRES in 40S/IRES_wt_ binary complexes is incompatible with binding of eIF2 and the closed conformation of the 40S subunit, and consequently, in 48S ICs, domain II is oriented away from the subunit interface, toward the solvent side, occupying multiple conformations oriented away from the E site. In eIF5B-containing pre-48S ICs, the 40S subunit is in the open conformation and domains II is inserted into the E site. Upon codon-anticodon base-pairing in eIF5B-containing 48S ICs, the 40S subunit adopts the closed conformation, and domain II becomes displaced from the E site and is oriented away from the subunit interface like in eIF2- containing 48S ICs. Importantly, compared to eIF2-containing 48S ICs, in eIF5B-containing pre-48S ICs and 48S ICs, Met-tRNAi^Met^ rotates by ∼14° and moves 15 Å from the head of the 40S subunit to a position that matches the orientation in 80S ribosomes. Thus, our data show how eIF5B repositions tRNA already on the 48S complex, thus preparing it for joining with the 60S subunit.

## Acknowledgments

This work was supported by NIH R01 GM55440 (to J.F.), R35 GM139453 (to J.F.), R35 GM122602 (to T.V.P.), and R01 AI123406 (to C.U.T.H.) Cryo-EM data were collected at the Columbia University Cryo-Electron Microscopy Center.

## Author Contributions

Conceptualization, T.V.P., C.U.T.H., and J.F.; Investigation, Z.P.B., and I.A.; Validation Z.P.B., I.A., and S.D.; Writing – Original Draft, T.V.P. and Z.P.B.; Writing – Review and Editing, Z.P.B., I.A., S.D., C.U.T.H., T.V.P., and J.F.; Funding Acquisition, C.U.T.H., T.V.P., and J.F.; Supervision, C.U.T.H., T.V.P., and J.F.

### Declaration of Interests

Authors declare no competing interests.

### SUPPLEMENTARY FIGURE LEGENDS

**Figure S1. Classification of HCV IRES containing ribosomal complexes, related to Figure 1.**

(A-D) Example micrographs for (A) the *wt* eIF2-containing sample, (B) the *wt* IRES eIF5B- containing sample, (C) the ΔdII IRES eIF2-containing sample, and (D) the ΔdII IRES eIF5B- containing sample. Further details in **Table S2**. Scale bar 2000 Å.

(E) Example 2D classification results for the *wt* IRES eIF2-containing sample.

(F-G) Classification scheme for (F) *wt* IRES and (G) ΔdII IRES containing samples. For full details regarding classification see methods. Final maps are colored showing the 40S subunit (yellow), IRES (red), eIF1A (blue), Met-tRNAi^Met^ (magenta), and initiation factors eIF2 or eIF5B (green). Full details for complexes are available in Table S3 (structures 1_ΔdII_-15_wt/ΔdII_), Table S5 (16_wt_-28_wt_), and Table S7 (structures 29_wt_-46_ΔdII_).

**Figure S2. Local resolution, related to Figure 1**.

(A) Local resolution of labeled HCV IRES complexes.

(B) Central slice of each complex.

Local resolution values and filtered maps can be found under EMDB ascension codes for each structure (see Table S3).

**Figure S3. Example densities, related to Figure 1**.

**(A)** Domain IIId (red) and ES7 (yellow) for the pre-48S IC (structure 14_wt_).

**(B)** Domain IIId (red) and ES7 (yellow) for the minimally bound complex (structure 1_ΔdII_)

**(C)** Contact between domain IIIa/IIIc (red) and riboprotein eS27 (magenta) in the ternary complex (structure 12_wt_).

**(D)** eIF2 alpha helix 1 in the *wt* IRES 48S IC (structure 12_wt_), top, and the ΔdII IRES 48S IC (structure 12_ΔdII_), bottom.

**(E)** eIF5B alpha helix 12 in the eIF5B-containing pre-48S IC (structure 14_wt_), top, and the 48S IC (structure 15_wt_), bottom.

**(F)** eIF1A example densities from the eIF5B-containing pre-48S IC (structure 14_wt_), left, and the eIF2-containing 48S IC (structure 12_wt_), right.

**(G)** Initiation codon (red) and tRNA anticodon (magenta) for all complexes where mRNA is present in the P site. Diagram shows identity of nucleotides and hydrogen bonds for tRNA and initiation codon bases in each accompanying figure.

**Figure S4. Interactions between HCV IRES and 40S ribosomal subunit for all complexes, related to Figures 1 and 2**.

Contacts between ribosomal proteins (red), 18S rRNA (yellow), and the AUG codon (blue) are marked. Labels refer to ribosomal proteins and elements of 18S rRNA, and to domains of the IRES. Complexes assembled using the ΔdII IRES show the deleted domain marked in red. Also see Table S4 for more details.

**Figure S5. eIF2 related supplement, related to Figure 4**.

**(A)** Overview of the ΔdII IRES eIF2-containing 48S initiation complex (structure 12_ΔdII_).

**(B-C)** Presence of Domain II in the E site causes the head of the 40S subunit to open. (B) Position of 18S rRNA in the open ternary complex (structure 11_wt_) (orange), *wt* IRES eIF2-containing 48S IC (structure 12_wt_) (yellow), and ΔdII IRES eIF2-containing 48S IC (structure 12_ΔdII_) (magenta).

(C) Position of 18S rRNA in the open ternary complex (structure 11_wt_) (orange), *wt* IRES w/o eIF2 48S IC (structure 13_wt_) (yellow), and ΔdII IRES w/o eIF2 48S IC (structure 13_ΔdII_) (magenta). Measured angle between 18S rRNA h33 and h28 in the ternary complex (denoted with ʹ) to h33 in the *wt* IRES complex (denoted with ʹʹ) shown for (C-D).

(D) Unsharpened maps for the *wt* IRES eIF2-containing 48S IC (structure 12_wt_), left, and the ΔdII IRES eIF2-containing 48S IC (structure 12_ΔdII_), right. Comparison to canonical cap-dependent initiation complexes (PDB: 6YAL) identifies the presence of eIF2γ (cyan).

(E) The position and conformation of mRNA (magenta) and the eIF2 α-subunit for the *wt* IRES eIF2-containing 48S IC (structure 12_wt_, green) and the ΔdII IRES eIF2-containing 48S IC (structure 12_ΔdII_ (orange) is identical.

(F) Maps showing that domain II occupies multiple positions in the *wt* IRES eIF2-containing 48S IC.

**(G-J)** Diagram of P site for the (G) IRES binary complex (structure 6_ΔdII_), **(H)** P site for pre-48S IC (structure 14_wt_), **(I)** P site for eIF2-containing 48S IC (structure 12_wt_), and **(J)** P site for eIF5B- containing 48S IC (structure 15_wt_). Ribosomal proteins and 18S rRNA (yellow), mRNA (red), tRNA (white), and eIF2 (green) are shown.

**Figure S6, related to Figure 5**.

**(A)** Overview of the ΔdII IRES eIF5B-containing 48S initiation complex (structure 15_ΔdII_).

**(B)** Maps showing that domain II occupies multiple positions in the *wt* IRES eIF5B-containing 48S IC.

**(C)** Position of 18S rRNA in the *wt* IRES pre-48S IC (structure 14_wt_) (orange), *wt* IRES eIF5B- containing 48S IC (structure 15_wt_) (magenta), and ΔdII IRES eIF5B-containing 48S IC (structure 15_ΔdII_) (yellow). Measured angle between 18S rRNA h33 and h28 in the eIF5B-containing 48S IC (denoted with ʹ) to h33 in the pre-48S IC (denoted with ʹʹ).

**(D)** Global position of eIF5B bound to the intersubunit face of the 40S ribosomal subunit. Interactions between 18S rRNA (yellow), uS12 (salmon), and eIF1A (blue) are shown.

**(E)** Coulomb potential for the GTP nucleotide in eIF5B G domain for pre-48S IC (structure 14_wt_), left, and 48S IC (structure 15_wt_), right.

**(F)** Comparison of conformation of switch 1 in GTP (orange) and GDP (blue), and switch 2 in GTP (magenta) and GDP (yellow) conformations from the *C. thermophilum* G domain (grey) (PDB: 4NCN). Conformational changes in domain II seen in the *O. cuniculus* 48S IC state (green) and the *S. cerevisiae* pre-elongation 80S state (red) (PDB: 6WOO).

**(G)** Domain alignment between *O. cuniculus* pre-48S IC (structure 14_wt_)/48S IC (structure 15_wt_, green) and *S. cerevisiae* eIF5B from the pre-elongation 80S (red) (PDB: 6WOO). Position of G domain switch 1 (magenta) and switch 2 (orange) are marked

**Figure S7, related to Figures 5 and 6**.

**(A)** Structure of the *O. cuniculus* eIF5B G domain (PDB: 4UJD) bound to the pre-elongation 80S ribosome (yellow) compared to the *O. cuniculus* pre-48S IC (structure 14_wt_, green).

**(B)** Structure of the *O. cuniculus* eIF5B G domain in the pre-48S IC (structure 14_wt_) (green).

**(C)** Structure of the *C. thermophilum* eIF5B G domain (PDB: 4NCN) (blue) compared to the *O. cuniculus* eIF5B G domain bound to the pre-48S IC (structure 14_wt_) (green).

**(D)** Structure of the *S. cerevisiae* eIF5B G domain (PDB: 6WOO) bound to the pre-elongation 80S ribosome (red) compared to the *O. cuniculus* pre-48S IC (structure 14_wt_) (green).

**(E-F)** Contacts between eS28 (salmon) and the HCV IRES (red) in the (E) pre-48S IC (structure 14_wt_) and (F) 48S IC (structure 15_wt_).

**(G)** Distance between P_OUT_ tRNA (PDB: 7A09) and the initiation codon in ternary complex (structure 11_wt_)

**(H)** Contacts between tRNA and the P site for the eIF2-containing 48S IC.

**Table S1. Sample composition, related to Figure 1**.

**Table S2. Data collection statistics, related to Figure 1**.

**Table S3. Refinement and validation statistics, related to Figure 1.**

**Table S4. Contacts between the HCV IRES and 40S subunit, related to Figure 2.** Distances below 3.5 Å are marked as ‘yes’, between 3.5-4.5 Å are marked as ‘weak’, and above 4.5 Å marked as ‘no’.

**Table S5. Maps showing HCV IRES domain II movement, related to Figures 4 and 5**.

**Table S6. Contacts between eIF5B and the 40S subunit, related to Figure 5**.

**Table S7. Additional maps produced during data processing, related to Figure 1 and S1.**

## STAR Methods

### Resource availability

### Materials availability

Requests for materials and additional information can be directed to Dr. Tatyana Pestova (;) or Dr. Joachim Frank.

### Data availability

Primary models and maps (Table S3) reported in this study were deposited in the Protein Data Bank (PDB) and Electron Microscopy Data Bank (EMDB) under the following accession codes: structure 1_ΔdII_ (PDB: XXXX; EMD-XXXX), structure 2_ΔdII_ (PDB: XXXX; EMD-XXXX), structure 3_ΔdII_ (PDB: XXXX; EMD-XXXX), structure 4_ΔdII_ (PDB: XXXX; EMD-XXXX), structure 5_ΔdII_ (PDB: XXXX; EMD-XXXX), structure 6_ΔdII_ (PDB: XXXX; EMD-XXXX), structure 7_ΔdII_ (PDB: XXXX; EMD-XXXX), structure 8_ΔdII_ (PDB: XXXX; EMD-XXXX), structure 9_ΔdII_ (PDB: XXXX; EMD-XXXX), structure 10_wt_ (PDB: XXXX; EMD-XXXX), structure 11_wt_ (PDB: XXXX; EMD-XXXX), structure 12_wt_ (PDB: XXXX; EMD-XXXX), structure 12_ΔdII_ (PDB: XXXX; EMD-XXXX), structure 13_wt_ (PDB: XXXX; EMD-XXXX), structure 13_ΔdII_ (PDB: XXXX; EMD-XXXX), structure 14_wt_ (PDB: XXXX; EMD-XXXX), structure 15_wt_ (PDB: XXXX; EMD-XXXX), and structure 15_ΔdII_ (PDB: XXXX; EMD-XXXX). For each entry half maps, the mask used for post processing, Fourier correlation curve, and local resolution maps have been deposited as additional files.

Additional maps (table S5) showing the movement of HCV IRES domain II were deposited in the EMDB under the following accession codes: structure 16_wt_ (EMD-XXXX), structure 17_wt_ (EMD-XXXX), structure 18_wt_ (EMD-XXXX), structure 19_wt_ (EMD-XXXX), structure 20_wt_ (EMD-XXXX), and structure 21_wt_ (EMD-XXXX), structure 22_wt_ (EMD-XXXX), structure 23_wt_ (EMD-XXXX), structure 24_wt_ (EMD-XXXX), structure 25_wt_ (EMD-XXXX), structure 26_wt_ (EMD-25555), structure 27_wt_ (EMD-XXXX), and structure 28_wt_ (EMD-XXXX). For each entry half maps, the mask used for post processing, and Fourier correlation curve have been deposited as additional files.

Maps obtained during data processing (Table S7) from particle stacks that were compositionally and conformationally identical that were later combined were deposited in the EMDB under the following accession codes: structure 29_wt_ (EMD-XXXX), structure 30_wt_ (EMD-XXXX), structure 31_ΔdII_ (EMD-XXXX), and structure 32_ΔdII_ (EMD-XXXX). For each entry half maps, the mask used for post processing, and Fourier correlation curve have been deposited as additional files.

Consensus maps (Table S7) that were used for focused classification were deposited in the EMDB under the following accession codes: structure 33_ΔdII_ (EMD-XXXX), structure 34_ΔdII_ (EMD-XXXX), structure 35_wt_ (EMD-XXXX), structure 36_wt_ (EMD-XXXX), structure 37_ΔdII_ (EMD-XXXX), structure 38_wt_ (EMD-XXXX), structure 39_wt_ (EMD-XXXX), and structure 40_ΔdII_ (EMD-XXXX). Other high-resolution maps obtained during classification were deposited in the EMDB under the following accession codes: structure 41_wt_ (EMD-XXXX), structure 42_wt_ (EMD-XXXX), structure 43_wt_ (EMD-XXXX), structure 44_wt_ (EMD-XXXX), structure 45_wt_ (EMD-XXXX), and structure 46_ΔdII_ (EMD-XXXX). For each entry half maps, the mask used for post processing, mask used for focused classification, and the Fourier correlation curve have been deposited as additional files.

### Method details

#### Plasmids

Vectors for expression of His_6_-tagged eIF1A (Pestova et al., 1998a) and *Escherichia coli* methionyl tRNA synthetase (Lomakin et al., 2006) have been described. The plasmid HCV-MSTN-Stop (Hashem et al., 2013) containing HCV Type 1 nt. 40–375 was used for transcription of mRNA containing the *wt* HCV IRES. A derivative for transcription of HCV IRES lacking domain II (containing HCV nt. 125-375) was made by GenScript Corp. (Piscataway, NJ). The HCV plasmids were linearized by BamHI, and mRNAs were transcribed using T7 RNA polymerase (Thermo Scientific).

#### Purification of factors, ribosomal subunits and aminoacylation of tRNA

Native mammalian 40S subunits, eIF2, eIF3 and eIF5B were purified from rabbit reticulocyte lysate (RRL) (Green Hectares), as described (Pisarev et al., 2007). Recombinant eIF1A and *Escherichia coli* methionyl tRNA synthetase were expressed and purified from *E. coli* as described (Pisarev et al., 2007).

For purification of native rabbit total tRNA, 200 ml RRL were centrifuged at 45,000 rpm for 4.5 h in a Beckman 50.2 Ti rotor at 4_o_C in order to pellet polysomes. The supernatant was dialyzed overnight against buffer A (20 mM Tris [pH 7.5], 4 mM MgCl_2_, 250 mM KCl, 2 mM DTT) and applied to a DE52 (Whatman) column equilibrated with buffer A. The tRNA was eluted with buffer B (20 mM Tris [pH 7.5], 3 mM MgCl_2_, 700 mM NaCl, 2 mM DTT) and precipitated overnight with 2.5 volumes of ethanol at -80_o_C. The precipitate was centrifuged at 13,000 rpm for 15 minutes and resuspended in 5 ml buffer C (100 mM Tris [pH 7.5], 5 mM MgCl_2_), phenol-chloroform (pH 4.7)) extracted and precipitated again with 0.3 M NaOAc and 2.5 volumes of ethanol. To isolate tRNA, the pellet was dissolved and subjected to gel filtration on a Superdex 75 column (Pestova and Hellen, 2001). Purified total tRNA was aminoacylated using *E. coli* methionyl tRNA synthetase (to obtain Met-tRNAi^Met^) as described (Pisarev et al., 2007).

### Assembly of ribosomal complexes

To form 48S initiation complexes, 7 pmol HCV IRES mRNA (*wt* or Δdomain ΙΙ mutant) were incubated with 3.5 pmol 40S subunits, 10 pmol eIF1A, 4.5 pmol eIF3, total native rabbit tRNA containing 3.5 pmol Met-tRNAi^Met^, and 6 pmol eIF2 or 10 pmol eIF5B in 40 μl buffer D (20 mM Tris [pH 7.5], 100 mM KAc, 2.5 mM MgCl_2_, 2 mM DTT, 0.25 mM spermidine, 1 mM ATP and 0.2 mM GTP) for 10 minutes at 37°C. The obtained complexes (containing 87.5 nM 40S subunits) were applied directly onto grids without dilution.

### Grid preparation and electron microscopy

Gold foil grids were prepared from Quantifoil gold mesh grids (Passmore and Russo, 2016). Initially, Quantifoil R0.6/1.0 300 mesh gold grids (Quantifoil Micro Tools GmbH) were visually inspected to check for uniformity and intactness of the Quantifoil layer and then placed into an Auto 306 Turbo Vacuum Coater (Edwards Vacuum) at a pressure of 10^3^ Pa and then gold wire (Ted Pella, Inc) was evaporated for approximately 8 minutes to create a 500 Å layer. Deposition thickness was determined using the inbuilt film thickness monitor. To remove the underlying Quantifoil carbon foil layer the grids were then treated with plasma using a Gatan Solarus 950 (Gatan Inc) operated at 25 W for 4 minutes with an argon/oxygen gas mixture.

To prepare hydrophilic grids, 30 minutes prior to sample application, grids were treated with plasma using a Gatan Solarus 950 (Gatan Inc) operated at 25W for 25 seconds with a hydrogen/oxygen gas mixture. These grids were then transferred to the environmental chamber of a Vitrobot Mark IV (Thermo Fisher Scientific) maintained at 4°C and 100% humidity. Here 3 μL of sample were applied and then blotted for 4 seconds with blot force 3 before immediate plunging into an a cooled (77K) ethane-propane mixture (Tivol et al., 2008) and then transferred to liquid nitrogen. Selected grids were screened to confirm sample composition and ice thickness using a Tecnai F20 electron microscope (Thermo Fisher Scientific) equipped with a field emission gun (FEG) operating at 200kV and a K2 summit direct electron detector (Gatan, Inc).

After screening grids from each sample, data collection was performed on a Tecnai Polara F30 (Thermo Fisher Scientific) equipped with an FEG operating at 300 kV and a K3 direct electron detector (Gatan, Inc). Movies were collected at a nominal magnification of 52,000× and defocus range of -0.5 to -2.5 μm in counting mode with a pixel size of 0.95 Å/pixel using the automated collection software Leginon (Potter et al., 1999; Carragher et al., 2000; Suloway et al., 2005). Each movie consisted of 40 frames recorded over 4 seconds with a total dose of 70.9 e^-^/Å^2^. Due to sample conditions the 40S ribosome may enter a preferred orientation and so potions of the data were collected with a 30° stage tilt (Table S2). For the *wt* IRES eIF2-containing sample 14,815 micrographs (14,815 at 30° stage tilt) were collected over 2 sessions, for the *wt* IRES eIF5B-containing sample 27,263 micrographs (20,509 at 30° stage tilt) were collected over 4 sessions, for the ΔdII IRES eIF2-containing sample 22,735 micrographs (17,695 at 30° stage tilt) were collected over 3 sessions, and for the ΔdII IRES eIF5B-containing sample 13,809 micrographs (13,809 at 30° stage tilt) were collected over 2 sessions (Table S2). Optical groups of micrographs with similar beam tilt values were identified using k-means clustering on the image shift values recorded by the microscope during data collection.

### Image processing

Gain references for each session were produced by visually screening ∼1000 micrographs to remove images that contained gold foil and then summing them using cisTEM *sum_all_tifs* (Grant et al., 2018). Movies were then aligned using MotionCor2 (Zheng et al., 2017) with dose weighting of 1.77 e^-^/Å^2^/frame and local patch correction with 8×5 patches. Initial CTF parameters were estimated using CTFFIND4 (Rohou and Grigorieff, 2015). Particle locations were identified using Topaz version 0.2.3 (Bepler et al. 2019) by initially downscaling all micrographs by 8× and then using the Topaz general model to identify particles. Particles with a confidence score below 0 were removed and the remaining positions rescaled for subsequent processing in Relion 3.1 (Scheres, 2012; 2016; Zivanov et al., 2018; 2019).

Initially, we identified 2,183,185 particles for the *wt* IRES eIF2-containing sample (147 particles per micrograph), 1,133,335 particles for the *wt* IRES eIF5B-containing sample (42 particles per micrograph), 2,213,826 particles for the ΔdII IRES eIF2-containing sample (97 particles per micrograph), and 1,459,506 particles for the ΔdII IRES eIF5B-containing sample (106 particles per micrograph). Particle locations were extracted from micrographs into downsampled boxes of 100×100 pixels (at 3.8 Å/pixel) to speed initial classification. This corresponds to 400×400-pixel boxes (at 0.95 Å/pixel) without downsampling. 25 iterations of 2D classification were performed to identify incorrectly picked particles, contamination, and other particles that were unable to be correctly aligned (e.g., due to poor SNR). Particles that were selected for removal were subjected to additional 2D classification to confirm that they did not contain clear 40S ribosome particles.

After initial screening of the 2D classification data the remaining particles for each sample were 1,201,923 particles for the *wt* IRES eIF2-containing sample (81 particles per micrograph), 736,700 particles for the *wt* IRES eIF5B-containing sample (36 particles per micrograph), 1,484,658 particles for the ΔdII IRES eIF2-containing sample (84 particles per micrograph), and 1,119,610 particles for the ΔdII IRES eIF5B-containing sample (81 particles per micrograph). For each sample, all particles were refined into a single model which was used to estimate the defocus values on a per-particle basis, followed by an additional refinement step, and then 3D classification without alignment into 10 classes for 25 iterations. This initial 3D classification was used to identify the major conformational states present in each sample, as well as further removal of poor-quality particles. For the *wt* IRES eIF2-containing sample we identified 580,938 particles in the closed state, 176,793 particles in the open sate, and removed 444,192 particles. For the *wt* IRES eIF5B-containing sample we identified 378,325 open state, 360,338 closed state, and removed 58,037 particles. For the ΔdII IRES eIF5B-containing sample we identified 883,893 particles in the closed state, and removed 62,455 particles. For the ΔdII IRES eIF2-containing sample we identified 615,125 particles in the closed state, 198,920 particles in the intermediate-open state, and removed 670,433 particles. All particles that were selected for removal were subjected to additional 2D classification and 3D refinement steps to confirm that they did not contain 40S ribosome complexes.

### General processing pathway

All particles were re-extracted at full-size (400×400-pixel box, 0.95 Å/pixel) and underwent iterations of 3D refinement, following by anisotropic magnification correction, defocus refinement, and beam tilt estimation. Multiple rounds of 3D classification (25 iterations, without alignment) were used to progressively remove poor quality particles. After each round of CTF refinement each particle stack underwent 3D refinement and was checked for increase in resolution and visual improvement of map density. Once CTF refinement no longer improved map quality, particle polishing using all frames was performed and then iterations of CTF refinement as outline above were completed. Focused classification on consensus maps was performed to isolate desired conformational or compositional states (see below).

### *wt* IRES eIF2-containing sample

After 3D classification the consensus map of closed 40S ribosome particles (Table S7) from the *wt* IRES eIF2-containing sample contained 530,720 particles at 3.1 Å resolution (structure 36_wt_; EMD-XXXX) underwent focused classification by enclosing Met-tRNAi^Met^ and the eIF2α subunit in a mask. Iterations of focused classification were able to produce two high quality classes of either the eIF2-containg 48S IC with 46,904 particles at 3.6 Å resolution (structure 12_wt_; PDB: XXXX; EMD-XXXX) and the 48S IC w/o eIF2 with 15,598 particles at 4.4 Å resolution (structure 13_wt_; PDB: XXXX; EMD-XXXX). An additional class of the eIF2-containg 48S IC lacking eIF1A with 45,571 particles at 3.6 Å resolution (structure 45_wt_; EMD-XXXX) was obtained. Extensive classification of the remaining 422,647 particles produced maps that showed the 40S ribosome bound to eIF2 and/or Met-tRNA_iMet_ but with a low resolution, or the ribosome lacking both factors.

To determine the positions that IRES domain II occupies the region around domain II was masked and classified into 10 classes over 25 iterations without alignment. Six classes were able to be clearly resolved showing IRES domain II sampling different positions around the ribosome head. Class 1 containing 41,393 particles at 3.8 Å resolution (structure 16_wt_; EMD-XXXX), class 2 containing 39,148 particles at 3.8 Å resolution (structure 17_wt_; EMD-XXXX), class 3 containing 49,034 particles at 3.7 Å resolution (structure 18_wt_; EMD-XXXX), class 4 containing 66,090 particles at 3.6 Å resolution (structure 19_wt_; EMD-XXXX), class 5 containing 24,190 particles at 3.9 Å resolution (structure 20_wt_; EMD-XXXX), and class 6 containing 36,155 particles at 3.8 Å resolution (structure 21_wt_; EMD-XXXX). Extensive classification of the remaining 274,710 particles could not further resolved additional positions of domain II.

After extracting at full size the consensus map of the open 40S ribosome particles (Table S7) from the *wt* IRES eIF2-containing sample contained 176,793 particles at 3.8 Å resolution (structure 29_wt_; EMD-XXXX).

### *wt* IRES eIF5-containing sample

The consensus map of closed 40S ribosome particles (Table S7) from the *wt* IRES eIF5B- containing sample contained 360,338 particles at 3.4 Å resolution (structure 39_wt_; EMD-XXXX) underwent focused classification by enclosing eIF5B in a mask. This could be classified into three high-resolution classes of the eIF5B-containing 48S IC eIF5B: one containing 133,782 particles at 3.7 Å resolution (structure 15_wt_; PDB: XXXX; EMD-XXXX), one containing 109,025 particles at 3.7 Å resolution (structure 41_wt_; EMD-XXXX), and one containing 55,367 at 3.8 Å resolution (structure 42_wt_; EMD-XXXX). The remaining 62,164 particles showed the 48S IC at 3.9 Å resolution but the occupancy of eIF5B was lower (structure 43_wt_; EMD-XXXX).

To determine the positions that IRES domain II occupies the region around domain II was masked and classified into 10 classes over 25 iterations without alignment. Seven classes were able to be resolved showing IRES domain II sampling different positions around the ribosome head. Class 1 containing 20,138 particles at 5.0 Å resolution (structure 22_wt_; EMD-XXXX), class 2 containing 11,082 particles at 5.4 Å resolution (structure 23_wt_; EMD-XXXX), class 3 containing 16,584 particles at 5.2 Å resolution (structure 24_wt_; EMD-XXXX), class 4 containing 16,217 particles at 4.9 Å resolution (structure 25_wt_; EMD-XXXX), class 5 containing 8,102 particles at 6.0 Å resolution (structure 26_wt_; EMD-XXXX), class 6 containing 13,152 particles at 5.3 Å resolution (structure 27_wt_; EMD-XXXX), and class 7 containing 7,029 particles at 6.2 Å resolution (structure 28_wt_; EMD-XXXX). Extensive classification of the remaining 268,034 particles could not further resolved the position of domain II.

The consensus map of open 40S ribosome particles (Table S7) from the *wt* IRES eIF5B- containing sample was classified into two classes: a consensus eIF5B-containing pre-48S IC with 199,047 particles at 3.6 Å (structure 38_wt_; EMD-XXXX) and the open binary complex with 147,309 particles at 4.1 Å resolution (structure 30_wt_; EMD-XXXX). The 31,969 remaining particles were subjected to 3D classification but did now contain high-quality classes. The consensus map of the eIF5B containing pre-48S IC (structure 38_wt_) was masked around eIF5B and focused classification produced two classes, one with 60,578 particles at 3.8 Å (structure 14_wt_; PDB: XXXX; EMD-XXXX), and a lower-resolution map from 29,072 particles at 4.2 Å (structure 44_wt_; EMD-XXXX).

### ΔdII IRES eIF2-containing sample

The consensus map of closed 40S ribosome particles (Table S7) from the ΔdII IRES eIF2- containing sample contained 615,195 particles at 3.3 Å resolution (structure 37_ΔdII_; EMD-25596) underwent focused classification by enclosing Met-tRNA_iMet_ and the eIF2α subunit in a mask. Iterations of focused classification were able to produce two high quality classes of either the eIF2- containg 48S IC (structure 12_ΔdII_; PDB: XXXX; EMD-XXXX), and the 48S IC w/o eIF2 (structure 13_ΔdII_; PDB: XXXX; EMD-XXXX). An additional class of 147,713 particles at 3.6 Å (structure 46_ΔdII_; EMD-XXXX) showed the 48S IC but with very poor density for eIF2. Extensive classification of the remaining 348,071 particles did not produced any high-quality maps. The consensus map of the intermediate conformation 40S ribosome particles (Table S7) from the ΔdII IRES eIF2-containing sample had 198,920 particles at 3.5 Å resolution (structure 31_ΔdII_; EMD-25590).

### ΔdII IRES eIF5B-containing sample

3D classification of the closed state eIF5B-containing 40S ribosome produced three classes that were processed further (Figure S1G; Table S7). The consensus map of closed 40S ribosome particles (Table S7) from the ΔdII IRES eIF5B-containing sample contained 148,763 particles at 3.4 Å resolution (structure 40_ΔdII_; EMD-XXXX) and underwent focused classification by enclosing eIF5B in a mask. Iterations of focused classification were able to produce one high quality class of the eIF5B-containg 48S IC (structure 15_ΔdII_; PDB: XXXX; EMD-XXXX).

Classification of the remaining 99,239 particles produced maps at either low resolution or lacking eIF5B. The consensus map showing the closed 40S ribosome with multiple HCV IRES states (Table S7) had 346,516 particles at 3.6 Å resolution (structure 33_ΔdII_; EMD-XXXX). A mask was prepared around IRES domains S1/S2/IIIe/IIIf and focused classification resolved six classes of the IRES in various association/dissociation states: (1) the early stage association between the IRES and 40S/IRES_ΔdII_ binary complex with 42,271 particles at 4.3 Å resolution (structure 1_ΔdII_; PDB: XXXX; EMD-XXXX), (2) the early stage association between the IRES and 40S/IRES_ΔdII_ binary complex with 28,684 particles at 4.6 Å resolution (structure 2_ΔdII_; PDB: 7SYH; EMD-XXXX), (3) the early stage association between the IRES and 40S/IRES_ΔdII_ binary complex with 24,545 particles at 4.5 Å resolution (structure 3_ΔdII_; PDB: XXXX; EMD-XXXX), (4) the early stage association between the IRES and 40S/IRES_ΔdII_ binary complex with 27,043 particles at 4.8 Å resolution (structure 4_ΔdII_; PDB: XXXX; EMD-XXXX), (5) the early stage association between the IRES and 40S/IRES_ΔdII_ binary complex with 48,757 particles at 4.2 Å resolution (structure 5_ΔdII_; PDB: XXXX; EMD-XXXX), and (6) the canonically bound IRES 40S/IRES_ΔdII_ binary complex with 29,657 particles at 4.5 Å resolution (structure 6_ΔdII_; PDB: XXXX; EMD-XXXX). The remaining 145,559 particles were classified into either low-resolution maps or with the 40S ribosome lacking the HCV IRES. The consensus map of the intermediate 40S ribosome particles (Table S7) from the ΔdII IRES eIF5B-containing sample had 287,087 particles at 4.5 Å resolution (structure 32_ΔdII_; EMD-XXXX).

### Combining ΔdII IRES intermediate 40S ribosome particles and focused classification

Both eIF2-, and eIF5B-containing samples prepared on the ΔdII IRES yielded classes that showed the HCV IRES bound to the 40S ribosome where the head of the ribosome was in multiple states (structure 31_ΔdII_-32_ΔdII_). Both maps were visually inspected and showed high similarity (unsharpened maps a correlation of 0.9834, and after B-factor sharpening 0.9295). We combined these particles into a consensus map of 456,311 particles at 3.5 Å resolution (structure 34_ΔdII_; EMD-XXXX) that underwent focused classification of the entire head region of the ribosome that produced three classes: the closed 40S/IRES_ΔdII_ binary complex with 59,660 particles at 4.8 Å resolution (structure 7_ΔdII_; PDB: XXXX; EMD-XXXX), the intermediate-open 40S/IRES_ΔdII_ binary complex with 144,252 particles at 4.0 Å resolution (structure 8_ΔdII_; PDB: XXXX; EMD-XXXX), and the open 40S/IRES_ΔdII_ binary complex with 46,095 particles at 4.6 Å resolution (structure 9_ΔdII_; PDB: XXXX; EMD-XXXX). The remaining 206,304 particles were classified into low resolution maps.

### Combining *wt* IRES open 40S ribosome particles and focused classification

Both eIF2-, and eIF5B-containing samples prepared on the *wt* IRES yielded classes that showed the HCV IRES bound to the 40S ribosome with IRES domain II inserted into the E site (structure 29_wt_-30_wt_). Both maps were visually inspected and showed high similarity (unsharpened maps correlation of 0.9945, and after B-factor sharpening 0.9464). We combined these particles into a consensus map of 324,102 particles at 3.3 Å resolution (structure 35_wt_; EMD-XXXX) that underwent focused classification of eIF1A that produced two classes: the binary complex with 119,320 particles at 3.8 Å resolution (structure 10_wt_; PDB: XXXX; EMD-XXXX), and the ternary complex with 204,782 particles at 3.8 Å resolution (structure 11_wt_; PDB: XXXX; EMD-XXXX).

### Model building, and refinement

For all data, where applicable, we were able to unambiguously fit the head and body of the 40S (PDB: 6D9J; Pisareva et al. 2018), HCV IRES (PDB: 5FLX; Yamamoto et al. 2015), eIF1A (PDB: 4KZZ; Lomakin and Steitz 2013), tRNA (PDB: 5K0Y; Simonetti et al. 2016) eIF2α subunit (PDB: 6O85; Kenner et al. 2019), and eIF5B (PDB: 4UJD; Yamamoto et al. 2014). Initial model fitting was performed using UCSF Chimera v1.14 (Pettersen et al., 2004) with additional modelling in Coot (Emsley and Cowtan, 2004). For regions of eIF5B that did not have available models (e.g., Switch 1) model building was performed independently and then cross-checked for consistency. All models underwent one round of Phenix geometry minimization and multiple rounds of PHENIX real-space refinement (Adams et al., 2010; Afonine et al., 2018).

### Figures

All figures were prepared using UCSF Chimera v1.14 (Pettersen et al., 2004).

### Quantification and statistical analysis

Global resolution estimates were calculated using the 0.143 FSC criterion (Rosenthal and Henderson, 2003). Local resolution maps were calculated using Relion 3.1 (Scheres, 2012; 2016; Zivanov et al., 2018; 2019) using the B-factor determined during post processing and the Modulation transfer function (MTF) curve for the K3 camera at 300 kV provided by the manufacturer. (https://www.gatan.com/techniques/cryo-em#MTF). RMSD calculations for 18S rRNA chains were performed using Pymol (Schrödinger, 2015). Model validation for all models were calculated using PHENIX (Adams et al., 2010; Afonine et al., 2018) installed as part of the SBGrid package (Morin et al., 2013).

## Notes

### Competing Interest Statement

The authors have declared no competing interest.

