## Supplementary figures and images for "Molecular architecture of 40S initiation complexes on the Hepatitis C virus IRES: from ribosomal attachment to eIF5B-mediated reorientation of initiator tRNA"

### Supplemental figures

Brown et al., Figure S1

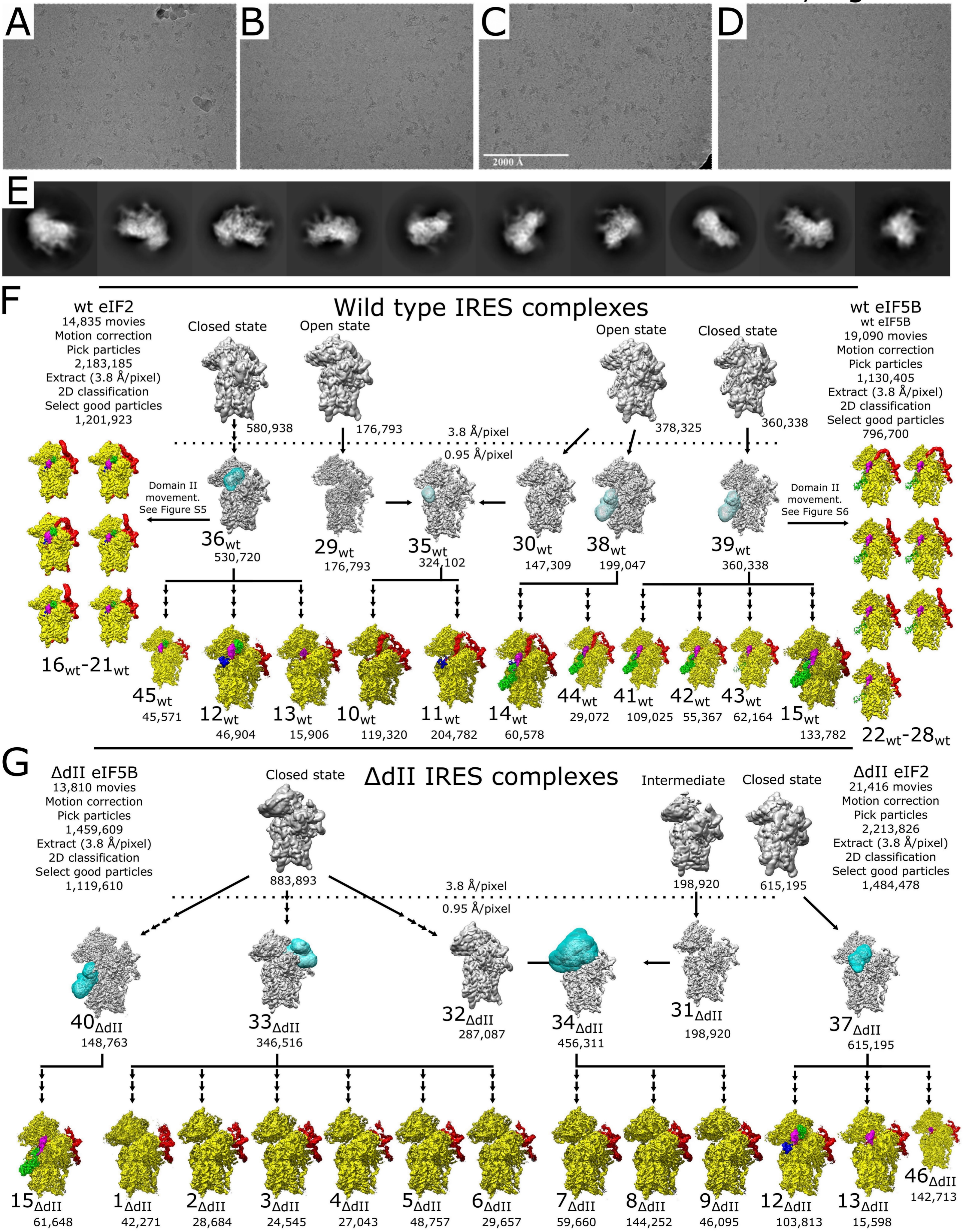

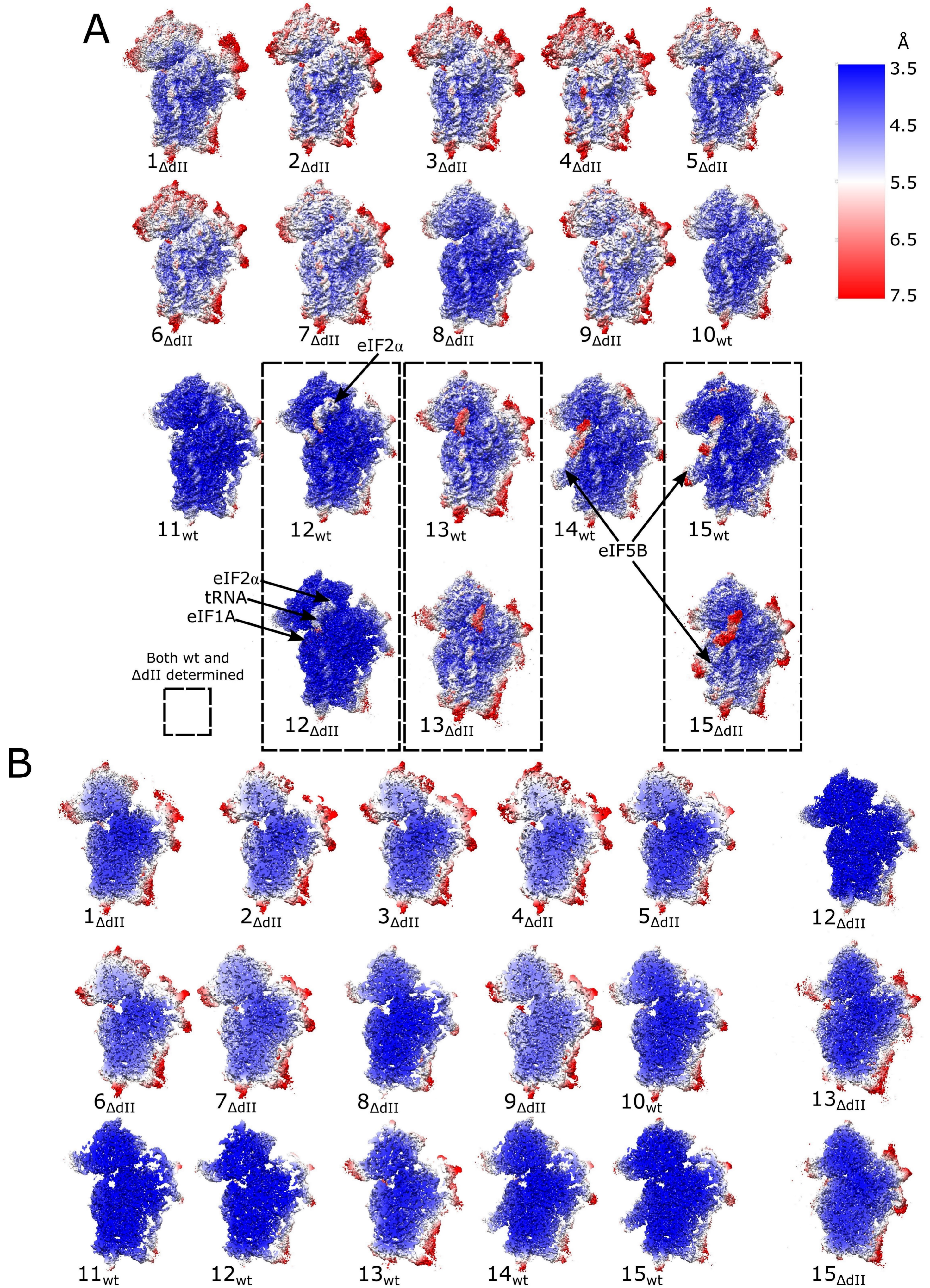

Brown et al., Figure S3

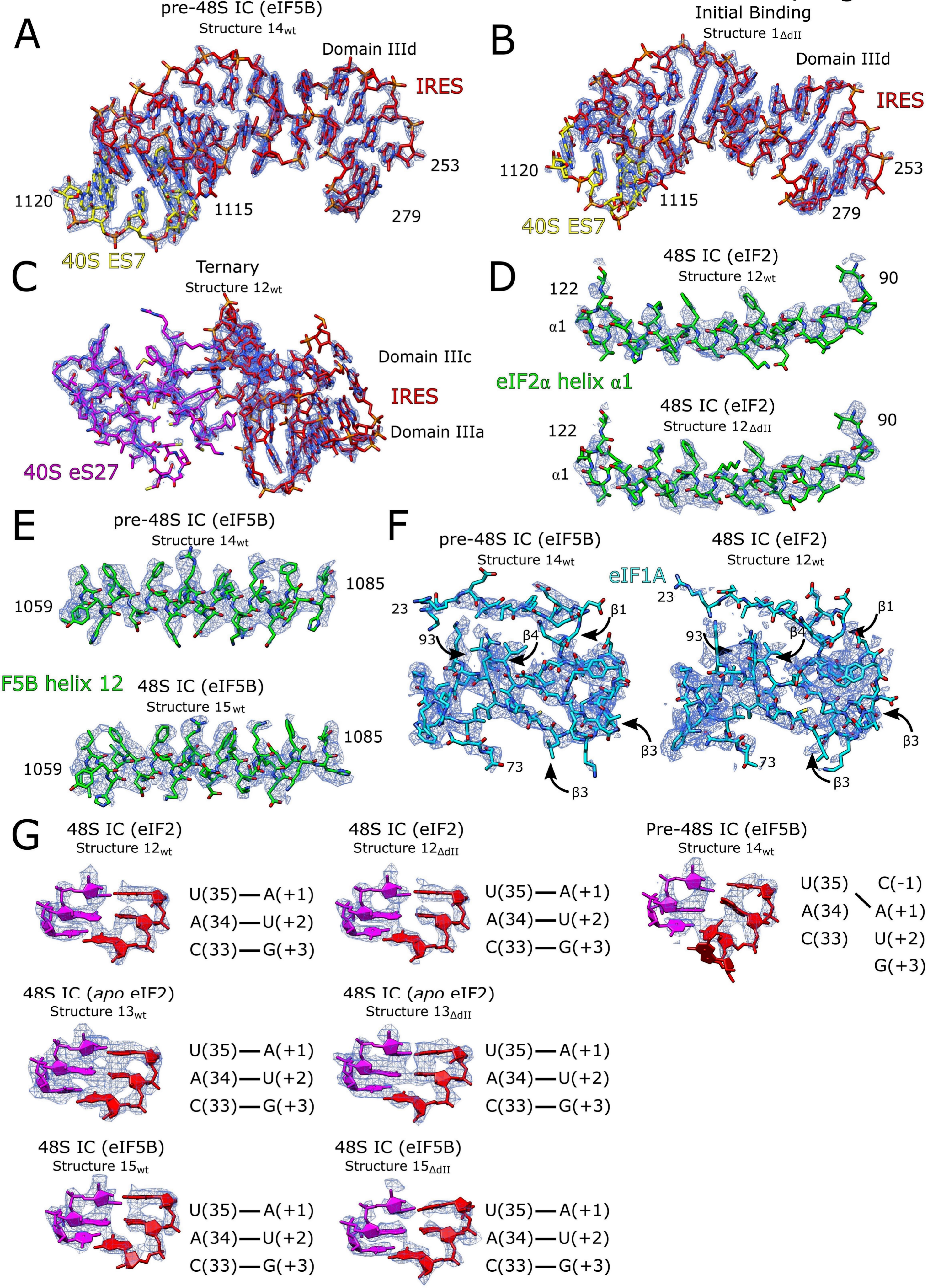

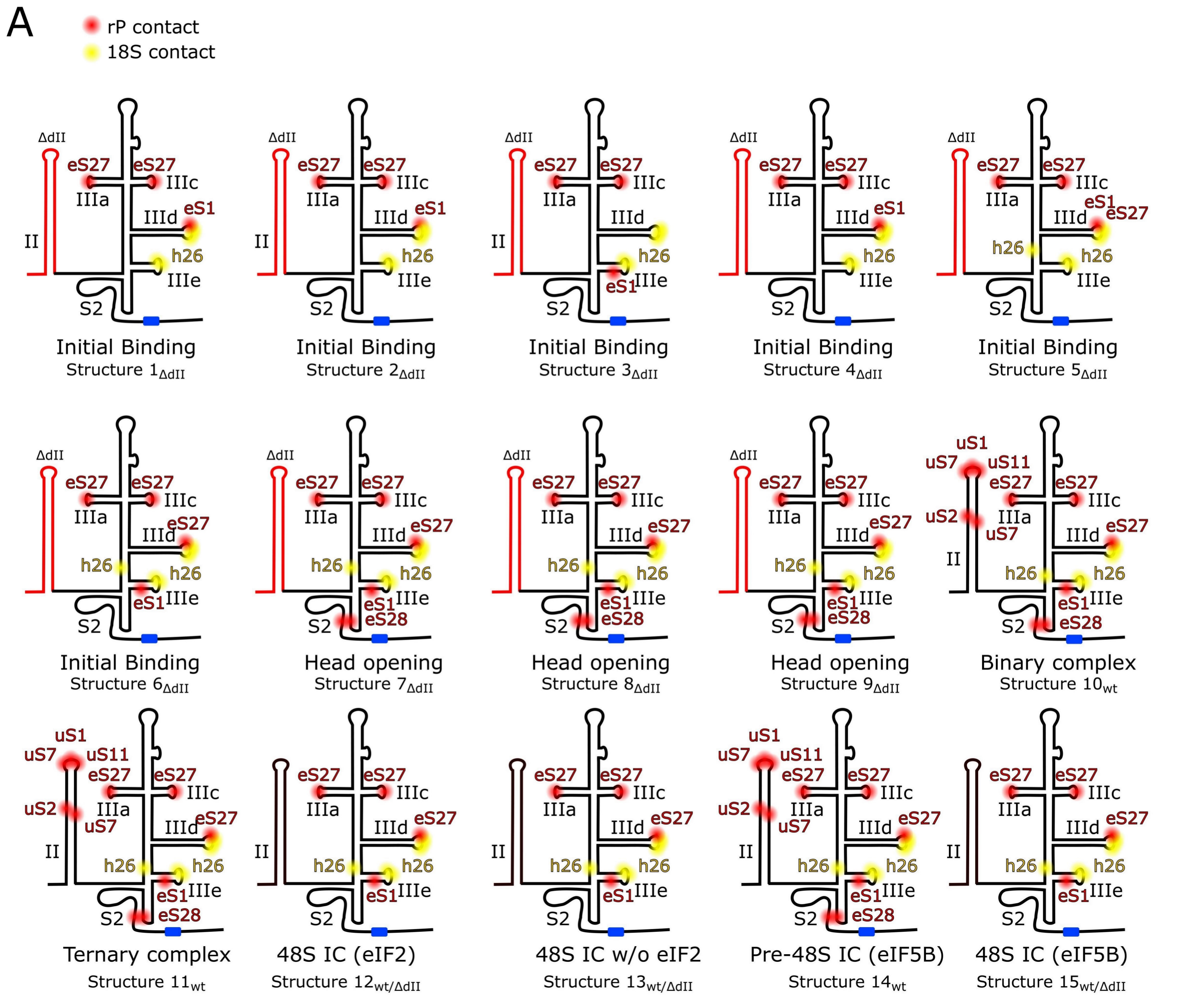

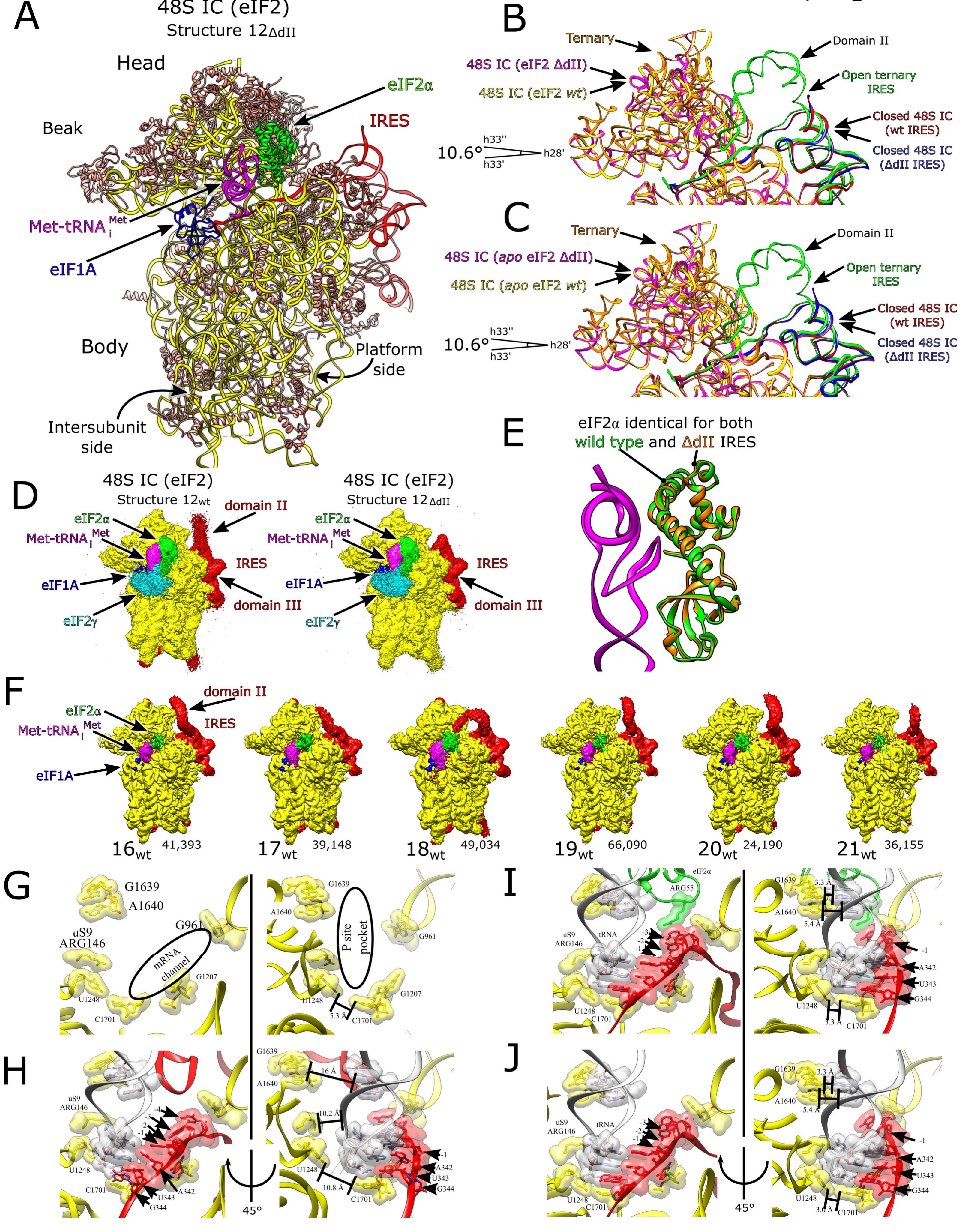

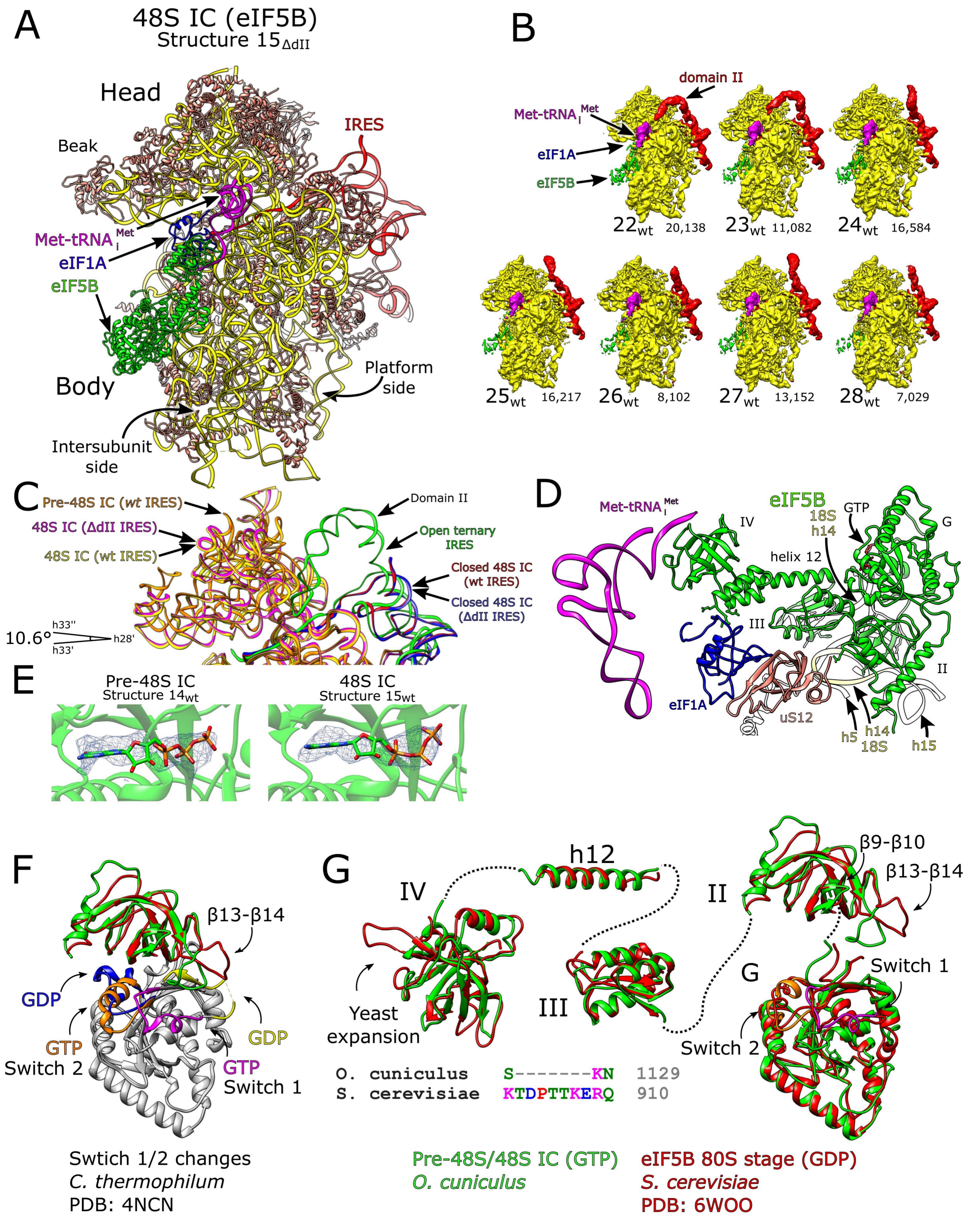

Brown et al., Figure S7

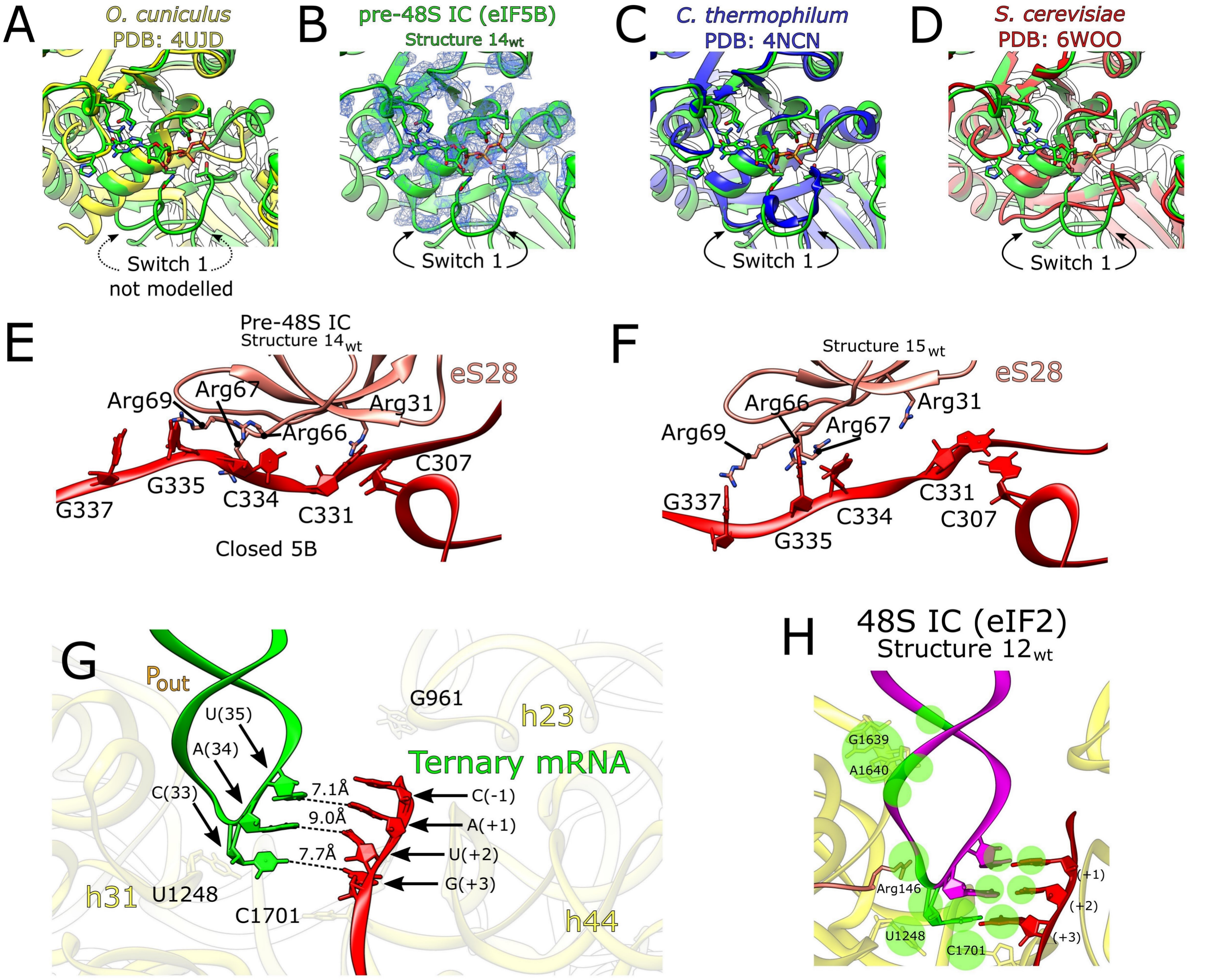
