## Supplementary material for "Molecular architecture of 40S initiation complexes on the Hepatitis C virus IRES: from ribosomal attachment to eIF5B-mediated reorientation of initiator tRNA": Table S1

**Table S1**. **Sample composition, related to Figure 1.**

| IRES | Wild type | Wild type | ΔdII | ΔdII |
| --- | --- | --- | --- | --- |
| Ternary Complex | eIF2 | eIF5B | eIF2 | eIF5B |
| 40S small subunit | 3.5 pmol | 3.5 pmol | 3.5 pmol | 3.5 pmol |
| Met-tRNA_i_^Met^ | 3.5 pmol | 3.5 pmol | 3.5 pmol | 3.5 pmol |
| eIF2 | 6 pmol | - | 6 pmol | - |
| eIF5B | - | 10 pmol | - | 10 pmol |
| eIF3 | 4.5 pmol | 4.5 pmol | 4.5 pmol | 4.5 pmol |
| eIF1A | 10 pmol | 10 pmol | 10 pmol | 10 pmol |
| *wt* HCV IRES | 7 pmol | 7 pmol | - | - |
| ΔdII HCV IRES | - | - | 7 pmol | 7 pmol |
