## Supplementary material for "Molecular architecture of 40S initiation complexes on the Hepatitis C virus IRES: from ribosomal attachment to eIF5B-mediated reorientation of initiator tRNA": Table S2

**Table S2. Data collection statistics, related to Figure 1.**

| IRES | Wild type | Wild type | ΔdII | ΔdII |
| --- | --- | --- | --- | --- |
| Ternary Complex | eIF2 | eIF5B | eIF2 | eIF5B |
| Electron microscope | Polara F30 | Polara F30 | Polara F30 | Polara F30 |
| Magnification | 52,000 | 52,000 | 52,000 | 52,000 |
| Detector | K3 Summit | K3 Summit | K3 Summit | K3 Summit |
| Pixel size (Å) | 0.95 | 0.95 | 0.95 | 0.95 |
| Voltage (kV) | 300 | 300 | 300 | 300 |
| Electron exposure (e-/Å2) | 70.9 | 70.9 | 70.9 | 70.9 |
| Defocus range (μm) | 1-2.5 | 1-2.5 | 1-2.5 | 1-2.5 |
| Symmetry imposed | C1 | C1 | C1 | C1 |
| Total collection sessions | 2 | 4 | 3 | 2 |
| Total micrographs | 14,815 | 27,263 | 22,735 | 13,809 |
| No. collected at 30° tilt | 14,815 | 20,509 | 17,695 | 13,809 |
| Initial no. picked particles | 2,183,185 | 1,133,335 | 2,213,826 | 1,459,609 |
| Initial particles per micrograph | 147 | 42 | 97 | 106 |
| Particles after 2D classification | 1,201,923 | 736,700 | 1,484,658 | 1,119,610 |
| After 2D per micrograph | 81 | 36 | 84 | 81 |
