## Supplementary material for "Molecular architecture of 40S initiation complexes on the Hepatitis C virus IRES: from ribosomal attachment to eIF5B-mediated reorientation of initiator tRNA": Table S3

**Table S3. Refinement and validation statistics, related to Figure 1.**

| Structure | 1ΔdII | 2ΔdII | 3ΔdII | 4ΔdII | 5ΔdII | 6ΔdII | 7ΔdII | 8ΔdII | 9ΔdII | 10wt | 11wt | 12wt | 12ΔdII | 13wt | 13ΔdII | 14wt | 15wt | 15ΔdII |
| --- | --- | --- | --- | --- | --- | --- | --- | --- | --- | --- | --- | --- | --- | --- | --- | --- | --- | --- |
| IRES | ΔdII | ΔdII | ΔdII | ΔdII | ΔdII | ΔdII | ΔdII | ΔdII | ΔdII | wt | wt | wt | ΔdII | wt | ΔdII | wt | wt | ΔdII |
| name |  |  |  |  |  |  |  |  |  | Binary | Ternary | 48S IC (eIF2) | 48S IC (eIF2) | 48S IC (w/o eIF2) | 48S IC (w/o eIF2) | pre-48S IC (eIF5B) | 48S IC (eIF5B) | 48S IC (eIF5B) |
| EMD |  |  |  |  |  |  |  |  |  |  |  |  |  |  |  |  |  |  |
| PDB |  |  |  |  |  |  |  |  |  |  |  |  |  |  |  |  |  |  |
| **Reconstruction** |  |  |  |  |  |  |  |  |  |  |  |  |  |  |  |  |  |  |
| Particles | 42,271 | 28,684 | 24,545 | 27,043 | 48,757 | 29,657 | 59,660 | 144,252 | 46,095 | 119,320 | 204,782 | 46,904 | 103,813 | 15,598 | 15,906 | 60,578 | 133,782 | 49,524 |
| Map resolution (Å) | 4.3 | 4.6 | 4.5 | 4.8 | 4.2 | 4.5 | 4.8 | 4.0 | 4.6 | 3.8 | 3.8 | 3.6 | 3.5 | 4.4 | 4.6 | 3.8 | 3.7 | 3.7 |
| FSC threshold | 0.143 | 0.143 | 0.143 | 0.143 | 0.143 | 0.143 | 0.143 | 0.143 | 0.143 | 0.143 | 0.143 | 0.143 | 0.143 | 0.143 | 0.143 | 0.143 | 0.143 | 0.143 |
| Map resolution range | 3.65-22.06 | 3.74-27.01 | 3.74-23.97 | 3.88-35.75 | 3.64-19.89 | 3.77-22.88 | 3.89-31.38 | 3.47-21.68 | 4.10-22.32 | 3.44-24.85 | 3.31-21.92 | 3.14-19.71 | 3.00-26.78 | 3.64-30.36 | 3.56-32.65 | 3.42-19.24 | 3.27-50.00 | 3.26-17.65 |
| **Refinement** |  |  |  |  |  |  |  |  |  |  |  |  |  |  |  |  |  |  |
| Model resolution (Å) | 4.3 | 4.5 | 4.5 | 4.7 | 4.2 | 4.7 | 4.4 | 3.8 | 4.4 | 4.0 | 3.8 | 3.7 | 3.3 | 4.1 | 4.3 | 3.8 | 3.7 | 3.8 |
| FSC threshold | 0.5 | 0.5 | 0.5 | 0.5 | 0.5 | 0.5 | 0.5 | 0.5 | 0.5 | 0.5 | 0.5 | 0.5 | 0.5 | 0.5 | 0.5 | 0.5 | 0.5 | 0.5 |
| Map sharpening factor | -103.01 | -113.25 | -93.88 | -102.84 | -104.30 | -101.41 | -142.94 | -128.11 | -138.92 | -111.66 | -114.32 | -81.81 | -83.72 | -101.78 | -76.23 | -74.85 | -89.27 | -57.82 |
| *Model composition:* |  |  |  |  |  |  |  |  |  |  |  |  |  |  |  |  |  |  |
| Chains | 37 | 38 | 38 | 38 | 38 | 38 | 38 | 38 | 38 | 40 | 39 | 40 | 40 | 38 | 38 | 42 | 40 | 41 |
| Protein residues | 4891 | 4891 | 4891 | 4891 | 4891 | 4891 | 4891 | 4891 | 4891 | 4896 | 4997 | 5177 | 5177 | 4896 | 4896 | 5643 | 5621 | 5622 |
| Nucleotides | 1859 | 1859 | 1859 | 1859 | 1859 | 1859 | 1859 | 1859 | 1859 | 1944 | 1944 | 1960 | 1960 | 1960 | 1960 | 2019 | 1960 | 1960 |
| non-hydrogen atoms | 78761 | 78761 | 78761 | 78761 | 78761 | 78748 | 78761 | 78761 | 78761 | 80505 | 81427 | 83005 | 82981 | 80962 | 80962 | 87809 | 86381 | 86744 |
| Ligands (Zn, Mg, GTP) | 2, 0, 0 | 2, 0, 0 | 2, 0, 0 | 2, 0, 0 | 2, 0, 0 | 2, 0, 0 | 2, 0, 0 | 2, 0, 0 | 2, 0, 0 | 2, 4, 0 | 2, 0, 0 | 1, 0, 0 | 1, 0, 0 | 1, 0, 0 | 1, 0, 0 | 1, 2, 1 | 1, 1, 1 | 1, 1, 1 |
| *B factors (Å):* |  |  |  |  |  |  |  |  |  |  |  |  |  |  |  |  |  |  |
| Protein | 75.56 | 89.93 | 84.93 | 113.27 | 71.04 | 55.13 | 94.07 | 63.67 | 114.01 | 77.89 | 80 | 42.53 | 47.53 | 93.17 | 73.03 | 77.75 | 60.49 | 32.88 |
| RNA | 100.74 | 113.53 | 100.93 | 133.44 | 92.69 | 66.72 | 115.92 | 92.63 | 111.55 | 99.54 | 101.29 | 59.87 | 67.35 | 108.3 | 109.43 | 93.22 | 84.3 | 53.5 |
| *RMS deviations:* |  |  |  |  |  |  |  |  |  |  |  |  |  |  |  |  |  |  |
| Bond lengths (Å) | 0.004 | 0.003 | 0.003 | 0.004 | 0.004 | 0.004 | 0.003 | 0.005 | 0.005 | 0.003 | 0.004 | 0.007 | 0.006 | 0.005 | 0.008 | 0.006 | 0.003 | 0.002 |
| Bond angles (°) | 0.64 | 0.614 | 0.642 | 0.713 | 0.681 | 0.694 | 0.624 | 0.745 | 0.769 | 0.584 | 0.61 | 0.815 | 0.72 | 0.724 | 1.051 | 0.799 | 0.547 | 0.503 |
| *Validation:* |  |  |  |  |  |  |  |  |  |  |  |  |  |  |  |  |  |  |
| Molprobity score | 2.15 | 2.18 | 2.21 | 2.17 | 2.19 | 2.25 | 2.13 | 2.18 | 2.35 | 2.01 | 1.98 | 1.93 | 1.94 | 2.13 | 2.49 | 2.18 | 1.87 | 1.80 |
| Clashscore | 13.73 | 15.52 | 16.03 | 17.31 | 15.70 | 17.19 | 14.81 | 13.84 | 20.63 | 13.03 | 11.99 | 8.81 | 9.32 | 14.31 | 25.59 | 14.01 | 8.88 | 9.41 |
| Poor rotamers (%) | 0.43 | 0.28 | 0.14 | 0.02 | 0.05 | 0.07 | 0.14 | 0.05 | 0.28 | 0.05 | 0 | 0.34 | 0.39 | 0.12 | 0.05 | 0.11 | 0.06 | 0.02 |
| *Ramachandran plot:* |  |  |  |  |  |  |  |  |  |  |  |  |  |  |  |  |  |  |
| Disallowed (%) | 0.04 | 0.06 | 0 | 0.02 | 0.02 | 0.02 | 0 | 0.02 | 0.04 | 0.02 | 0.02 | 0.02 | 0.02 | 0 | 0 | 0.09 | 0.04 | 0 |
| Favoured (%) | 91.60 | 92.16 | 91.52 | 93.42 | 91.79 | 91.29 | 92.78 | 90.54 | 90.29 | 94.38 | 94.25 | 92.77 | 93.10 | 92.58 | 88.44 | 90.70 | 94.10 | 95.69 |
| Allowed (%) | 99.96 | 99.94 | 100 | 99.97 | 99.98 | 99.98 | 100 | 99.98 | 99.96 | 99.97 | 99.98 | 99.98 | 99.98 | 100 | 100 | 99.91 | 99.96 | 100 |
| *RNA validation:* |  |  |  |  |  |  |  |  |  |  |  |  |  |  |  |  |  |  |
| Length outliers (%) | 0.00 | 0.00 | 0.00 | 0.00 | 0.00 | 0.00 | 0.00 | 0.00 | 0.00 | 0.00 | 0.00 | 0.00 | 0.00 | 0.00 | 0.00 | 0.00 | 0.00 | 0.00 |
| Angle outliers (%) | 0.00 | 0.00 | 0.00 | 0.00 | 0.00 | 0.00 | 0.00 | 0.00 | 0.00 | 0.00 | 0.00 | 0.00 | 0.00 | 0.00 | 0.00 | 0.00 | 0.00 | 0.00 |
| Sugar pucker outliers (%) | 0.01 | 0.01 | 0.01 | 0.01 | 0.01 | 0.01 | 0.01 | 0.01 | 0.01 | 0.01 | 0.01 | 0.01 | 0.01 | 0.01 | 0.01 | 0.00 | 0.01 | 0.01 |
| Average suit | 0.468 | 0.468 | 0.456 | 0.446 | 0.468 | 0.416 | 0.450 | 0.445 | 0.416 | 0.501 | 0.503 | 0.480 | 0.511 | 0.475 | 0.365 | 0.460 | 0.544 | 0.545 |
