## Supplementary material for "Molecular architecture of 40S initiation complexes on the Hepatitis C virus IRES: from ribosomal attachment to eIF5B-mediated reorientation of initiator tRNA": Table S4

**Table S4. Contacts between the HCV IRES and 40S subunit, related to Figure 2.** Distances below 3.5 Å are marked as ‘yes’, between 3.5-4.5 Å are marked as ‘weak’, and above 4.5 Å marked as ‘no’.

| **IRES Domain** | **No.** | **nt** | **40S** | **No.** | **Res.** | **1ΔdII** | **2ΔdII** | **3ΔdII** | **4ΔdII** | **5ΔdII** | **6ΔdII** | **7ΔdII** | **8ΔdII** | **9ΔdII** | **10wt** | **11wt** | **12wt** | **12ΔdII** | **13wt** | **13ΔdII** | **14wt** | **15wt** | **15ΔdII** |
| --- | --- | --- | --- | --- | --- | --- | --- | --- | --- | --- | --- | --- | --- | --- | --- | --- | --- | --- | --- | --- | --- | --- | --- |
| II | 68 | G | eS25 | 78 | Lys | no | no | no | no | no | no | no | no | no | yes | yes | no | no | no | no | yes | no | no |
| II | 81 | A | uS7 | 130, 133 | Arg, Thr | no | no | no | no | no | no | no | no | no | yes | yes | no | no | no | no | yes | no | no |
| II | 83-86 | C | uS11 | 66 | Arg | no | no | no | no | no | no | no | no | no | yes | yes | no | no | no | no | yes | no | no |
| II | 101 | U | eS25 | 76 | Arg | no | no | no | no | no | no | no | no | no | yes | yes | no | no | no | no | yes | no | no |
| hIII1 | 136 | A | 18S | 1115 | U | no | no | no | no | yes | yes | yes | yes | yes | yes | yes | yes | yes | yes | yes | yes | yes | yes |
| IIIa | 163/164 | G/U | eS27 | 41 | Tyr | yes | yes | yes | yes | yes | yes | yes | yes | yes | yes | yes | yes | yes | yes | yes | yes | yes | yes |
| IIIc | 233 | G | eS27 | 43,80 | Ile,Arg | yes | yes | yes | yes | yes | yes | yes | yes | yes | yes | yes | yes | yes | yes | yes | yes | yes | yes |
| IIId | 264/265 | U | eS1 | 199 | Lys | yes | yes | no | yes | yes | weak | no | no | no | no | no | no | yes | no | no | yes | weak | weak |
| IIId | 266 | G | 18S | 1118 | C | no | yes | yes | yes | yes | yes | yes | yes | yes | yes | yes | yes | yes | yes | yes | yes | yes | yes |
| IIId | 267 | G | 18S | 1117 | C | no | yes | yes | yes | yes | yes | yes | yes | yes | yes | yes | yes | yes | yes | yes | yes | yes | yes |
| IIId | 266/267 | G/G | eS27 | 75 | Glu | no | weak | weak | yes | yes | yes | yes | yes | yes | no | no | yes | yes | no | no | weak | weak | yes |
| IIId | 268 | G | 18S | 1116 | C | no | no | yes | yes | yes | yes | yes | yes | yes | yes | yes | yes | yes | yes | yes | yes | yes | yes |
| IIIe | 295 | G | 18S | 1114 | U | no | yes | yes | yes | yes | yes | yes | yes | yes | yes | yes | yes | yes | yes | yes | yes | yes | yes |
| IIIe | 296 | A | 18S | 1114 | U | no | no | no | no | yes | yes | yes | yes | yes | yes | yes | yes | yes | yes | yes | yes | yes | yes |
| IIIe | 295 | G | 18S | 1115 | U | yes | no | no | no | no | no | no | no | no | no | no | no | no | no | no | no | no | no |
| IIIe | 295 | G | eS1 | 199 | Lys | no | weak | yes | no | no | no | weak | weak | no | weak | weak | yes | no | weak | yes | yes | no | no |
| IIIe | 296 | A | 18S | 1115 | U | no | yes | yes | no | no | no | no | no | no | no | no | no | no | no | no | no | no | no |
| IIIe | 300 | G | eS1 | 147 | Asn | no | no | no | no | yes | yes | yes | yes | yes | yes | yes | yes | yes | yes | yes | yes | yes | yes |
