## Supplementary material for "Molecular architecture of 40S initiation complexes on the Hepatitis C virus IRES: from ribosomal attachment to eIF5B-mediated reorientation of initiator tRNA": Table S5

**Table S5. Maps showing HCV IRES domain II movement, related to Figures 4 and 5.**

| Structure | 16_wt_ | 17_wt_ | 18_wt_ | 19_wt_ | 20_wt_ | 21_wt_ | 22_wt_ | *23_wt_* | *24_wt_* | 25_wt_ | 26_wt_ | 27_wt_ | 28_wt_ |
| --- | --- | --- | --- | --- | --- | --- | --- | --- | --- | --- | --- | --- | --- |
| Sample: | eIF2 | eIF2 | eIF2 | eIF2 | eIF2 | eIF2 | eIF5B | eIF5B | eIF5B | eIF5B | eIF5B | eIF5B | eIF5B |
| EMDB |  |  |  |  |  |  |  |  |  |  |  |  |  |
| **Reconstruction** |  |  |  |  |  |  |  |  |  |  |  |  |  |
| Particles | 41,393 | 39,148 | 49,034 | 66,090 | 24,190 | 36,155 | 20,138 | 11,082 | 16,584 | 16,217 | 8,102 | 13,152 | 7,029 |
| Map resolution (Å) | 3.8 | 3.8 | 3.7 | 3.6 | 3.9 | 3.8 | 5.0 | 5.4 | 5.2 | 4.9 | 6.0 | 5.3 | 6.2 |
| FSC threshold | 0.143 | 0.143 | 0.143 | 0.143 | 0.143 | 0.143 | 0.143 | 0.143 | 0.143 | 0.143 | 0.143 | 0.143 | 0.143 |
| Map sharpening factor | -79.90 | -90.02 | -71.72 | -86.21 | -93.54 | -82.41 | -169.46 | -144.09 | -156.72 | -120.45 | -135.74 | -108.45 | -132.07 |
| Map resolution range | 3.2-13.1 | 3.3-13.7 | 3.2-20.2 | 3.1-14.3 | 3.4-16.4 | 3.3-14.5 | 4.3-20.6 | 4.6-21.2 | 4.4-23.8 | 4.0-42.6 | 4.4-49.0 | 4.1-23.9 | 45.-26.3 |
