## Supplementary material for "Molecular architecture of 40S initiation complexes on the Hepatitis C virus IRES: from ribosomal attachment to eIF5B-mediated reorientation of initiator tRNA": Table S6

**Table S6. Contacts between eIF5B and the 40S subunit, related to Figure 5.** Contacts between eIF5B in the pre-48S IC (structure 14_wt_), 48S IC (structure 15_wt/ΔdII_), or yeast eIF5B-containing 80S (PDB: 6WOO).

| **Domain** | **no.** | **res.** | **chain** | **Helix** | **no.** | **res.** | **14wt** | **15wt** | **15ΔdII** | **PDB: 6WOO** |
| --- | --- | --- | --- | --- | --- | --- | --- | --- | --- | --- |
| II | 957 | GLU | uS12 | n/a | 142 | ARG | yes | no | yes | yes |
| II | 965 | ARG | 18S | 15 | 478 | G | yes | yes | yes | no |
| II | 965 | ARG | 18S | 15 | 488 | OP1 | yes | yes | yes | yes |
| II | 980 | ASN | 18S | 5 | 50 | A | yes | yes | yes | no |
| II | 984 | LYS | 18S | 5 | 488 | U | yes | yes | no | yes |
| II | 999 | LYS | 18S | 5 | 487 | U | yes | yes | yes | yes |
| III | 1077 | HIS | 18S | 5 | 480 | OP1 | yes | yes | yes | yes |
| III | 1078 | LYS | 18S | 5 | 482 | OP1 | no | no | no | no |
| III | 1103 | ARG | 18S | 14 | 460 | A | yes | yes | yes | no |
| III | 1103 | ARG | 18S | 14 | 461 | U | yes | yes | yes | no |
| III | 1106 | ARG | 18S | 14 | 459 | U | yes | yes | yes | no |
