## Supplementary material for "Molecular architecture of 40S initiation complexes on the Hepatitis C virus IRES: from ribosomal attachment to eIF5B-mediated reorientation of initiator tRNA": Table S7

**Table S7. Additional maps produced during data processing, related to Figure 1 and S1.**

| Structure | EMDB (PP job) | Particles | Map resolution (Å) | FSC threshold | Map sharpening factor | Note |
| --- | --- | --- | --- | --- | --- | --- |
| 29_wt_ |  | 176,793 | 3.8 | 0.143 | -132.09 | Pre-consensus (binary, eIF2) |
| 30_wt_ |  | 147,309 | 4.1 | 0.143 | -113.23 | Pre-consensus (binary, eIF5B) |
| 31_ΔdII_ |  | 198,920 | 3.5 | 0.143 | -85.69 | Pre-consensus (head movement, eIF2) |
| 32_ΔdII_ |  | 287,087 | 4.5 | 0.143 | -218.18 | Pre-consensus (head movement, eIF5B) |
| 33_ΔdII_ |  | 346,516 | 3.6 | 0.143 | -112.77 | Consensus (IRES binding) |
| 34_ΔdII_ |  | 456,311 | 3.5 | 0.143 | -104.49 | Consensus (head movement) |
| 35_wt_ |  | 342,102 | 3.3 | 0.143 | -99.38 | Consensus (binary) |
| 36_wt_ |  | 530,720 | 3.1 | 0.143 | -114.46 | Focused (eIF2) |
| 37_ΔdII_ |  | 615,195 | 3.3 | 0.143 | -108.92 | Focused (eIF2) |
| 38_wt_ |  | 199,047 | 3.6 | 0.143 | -84.45 | Focused (open, eIF5B) |
| 39_wt_ |  | 360,338 | 3.4 | 0.143 | -107.37 | Focused (closed, eIF5b) |
| 40_ΔdII_ |  | 148,763 | 3.4 | 0.143 | -65.49 | Focused (closed, eIF5B) |
| 41_wt_ |  | 109,025 | 3.7 | 0.143 | -97.61 | Closed, eIF5B |
| 42_wt_ |  | 55,367 | 3.8 | 0.143 | -89.55 | Closed, eIF5B |
| 43_wt_ |  | 62,164 | 3.9 | 0.143 | -117.14 | Closed, eIF5B |
| 44_wt_ |  | 29,072 | 4.2 | 0.143 | -70.09 | Open, eIF5B |
| 45_wt_ |  | 45,571 | 3.6 | 0.143 | -82.06 | Closed, eIF2 |
| 46_ΔdII_ |  | 142,713 | 3.6 | 0.143 | -104.37 | Closed, eIF2 ΔdII |
